# Integrative single cell multiomics analysis of human retina indicates a role for hierarchical transcription factors collaboration in genetic effects on gene regulation

**DOI:** 10.1101/2022.11.16.516814

**Authors:** Jun Wang, Xuesen Cheng, Qingnan Liang, Leah A. Owen, Meng Wang, Margaret M. DeAngelis, Yumei Li, Rui Chen

**Author notes:** Correspondence to be addressed to Rui Chen. These authors contributed equally (Jun Wang, Xuesen Cheng).

## Abstract

**Background:** Systematic characterization of how genetic variation modulates gene regulation in a cell type specific context is essential for understanding complex traits. To address this question, we profiled gene expression and chromatin state of cells from healthy retinae of 20 human donors with a single-cell multiomics approach, and performed genomic sequencing.

**Results:** We mapped single-cell eQTL (sc-eQTLs), single-cell caQTL (sc-caQTL), single-cell allelic specific chromatin accessibility (sc-ASCA) and single-cell allelic specific expression (sc-ASE) in major retinal cell types. By integrating these results, we identified and characterized regulatory elements and genetic variants effective on gene regulation in individual cell types. Most of the sc-eQTLs and sc-caQTLs identified show cell type specific effects, while the cis-elements containing the genetic variants with cell type specific effects tend to be accessible in multiple cell types. Furthermore, the transcription factors with binding sites perturbed by genetic variants tend to have higher expression in the cell types, where the variants have effect, than the cell types where the variants do not have effect. Finally, we identified the enriched cell types, candidate causal variants and genes, and cell type specific regulatory mechanism underlying GWAS loci.

**Conclusions:** Overall, genetic effects on gene regulation are highly context dependent. Our results suggest that among cell types sharing a similar lineage, cell type dependent genetic effect is primarily driven by trans-factors rather than cell type specific chromatin state of cis-elements. Our findings indicate a role for hierarchical transcription factors collaboration in cell type specific effects of genetic variants on gene regulation.

## Background

Gene regulation is cell type dependent[1], and the modulation of this process by genetic variation among individuals is a major contributor to complex traits and diseases [2–5]. Substantial progress has been made in mapping, annotation, and functional validation of regulatory variants[6–10]. However, the mechanisms by which genetic variants modulate gene regulation in cell type specific context remain largely unclear[11,12]. Indeed, prior *in vivo* studies conducted on bulk tissues have a limited ability to elucidate the cell type effects of gene regulation. This gap can be addressed by recent advances in single-cell omics technologies[1,13–16]. Recent studies using single-cell omics technologies, have generated cell atlases for different tissues and development stages, revealing regulatory elements in cell type/state resolution, facilitating the interpretation of non-coding variants[17–20]. Several pioneer studies further mapped expression QTL (eQTL) or chromatin accessibility QTL (caQTL) alone, based on molecular phenotypes profiled by single cell sequencing, which uncover the cell type/state specific effect of genetic variants[21–27]. Even so, the mechanisms underlying cell type/state specific effects of genetic variants are still elusive. To answer these questions, we integrated genomic sequencing with single cell multiomics profiling of gene expression and chromatin state, which offers a unique opportunity to identify and characterize regulatory elements, the effect of genetic variants, and the modulation mechanism underlying gene regulation in individual cell type contexts *in vivo*.

We performed whole genome sequencing (WGS), single nuclei RNA-sequencing (snRNA-seq) and single-nuclei assay for transposase-accessible chromatin sequencing (snATAC-seq) on the cells of healthy retinae from 20 human donors. We mapped sc-eQTLs, sc-caQTLs, sc-ASE, and sc-ASCA for major retinal cell types. Integration of these results leads to genome-wide identification and characterization of gene regulatory elements, and genetic variants affecting chromatin state and gene expression in individual cell type contexts. Intriguingly, most of sc-QTLs identified are specific to one cell type, suggesting a significant proportion of variants modulate gene expression and chromatin state depending on cell type. Further analyses suggest for the cell types sharing a similar lineage, such as retinal cell types studied here, the cell type specific effect of genetic variants seems not primarily due to cell type specific chromatin state of the affected cis-elements, but may be driven by perturbing the binding of trans-regulators. Finally, by integrating the single cell multiomics data, genetic association results and GWAS, we identified the enriched cell types, fine-mapped candidate causal variants and genes, and uncovered the regulatory mechanisms underlying GWAS loci.

## Results

### Single nuclei multiomics profiling of 20 healthy human donor retinae

To profile gene expression and chromatin state in cell type specific context, we performed snRNA-seq and snATAC-seq on the healthy retinae from 20 human donors (Fig. 1a, Supplementary Table 1). For snRNA-seq, upon quality control (QC), a total of 192,792 nuclei were clustered into 10 major retinal cell classes, including rod photoreceptors (Rod), cone photoreceptors (Cone), bipolar cells (BC), amacrine cells (AC), horizontal cells (HC), müller glia cells (MG), retinal ganglion cells (RGC), astrocytes (Astro), endothelial cells and microglia cells (Methods, Fig. 1b). In parallel, snATAC-seq was performed for the same set of donor retinae.

**Fig. 1:**
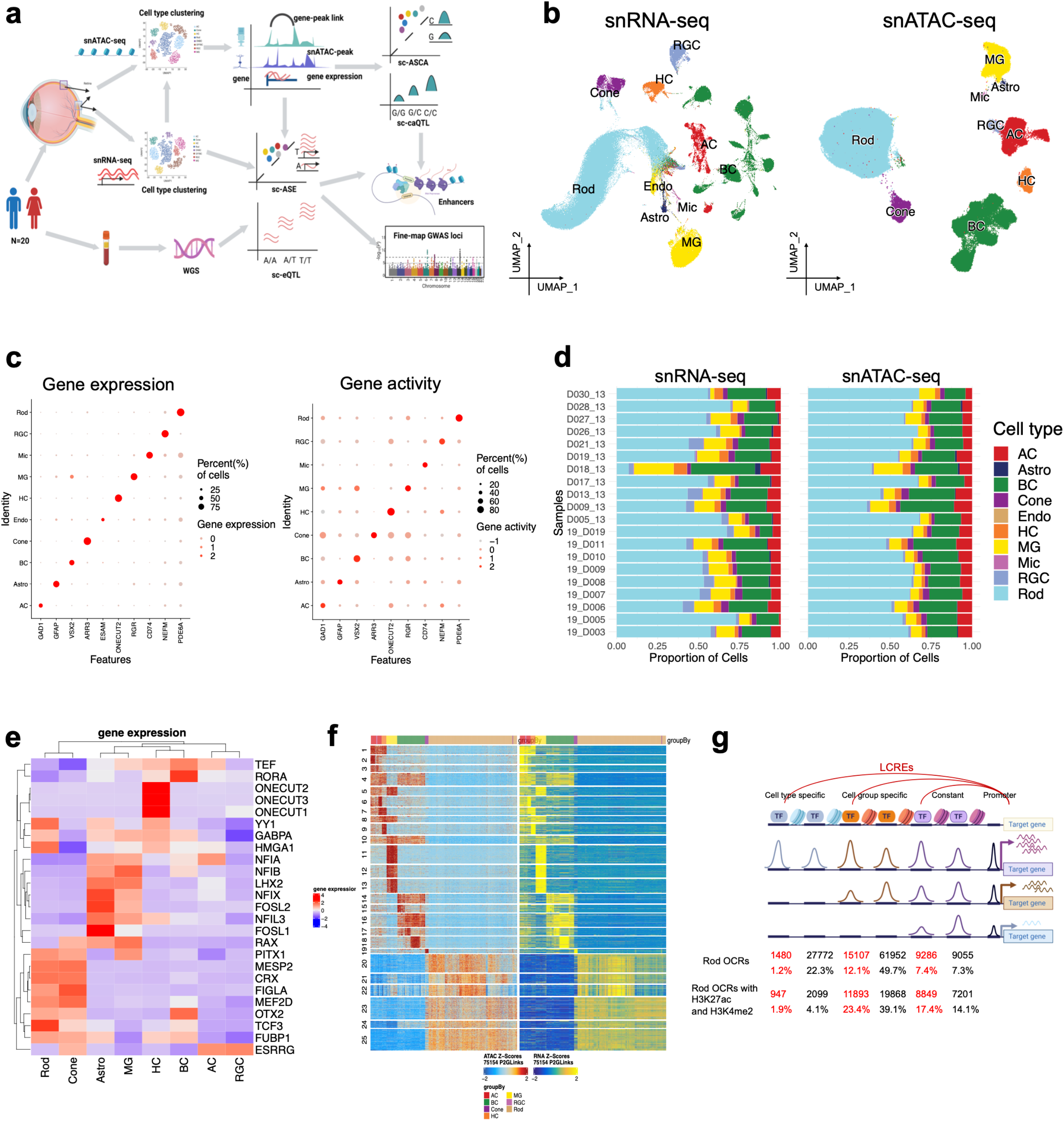
Profiling gene expression and chromatin accessibility of retinal cells. **a** Schematics of experiment design. **b** Uniform Manifold Approximation and Projection (UMAP) of cells from snRNA-seq and snATAC-seq. The cells were clustered into major retinal cell types. The same cell types from the two modalities are labeled with the same colors. **c** Marker gene expression measured by snRNA-seq and marker gene activity scores derived from chromatin accessibility measured by snATAC-seq are specific in the corresponding cell type. **d** The proportion of each cell type from snRNA-seq and snATAC-seq is similar across different samples. The number of cells per cell type per sample was listed in Supplementary Table 2. **e** Heatmap shows gene expression of the transcription factors identified in each cell type, based on chromVAR and the correlation between motif enrichment and gene expression. **f** Heatmap shows the chromatin accessibility (left) and gene expression (right) of 75154 significantly linked CRE-gene pairs. Rows were clustered using k-means clustering (k=25). **g** The proportions of Rod OCRs that are cell type specific LCRE, cell type specific non-LCRE, cell group specific LCRE, cell group specific non-LCRE, constant LCRE, and constant non-LCRE.

After QC, a total of 245,541 nuclei were clustered into 9 major retinal cell classes (Fig. 1b). Consistent with the cell type annotation, canonical cell type marker genes show specific expression and gene activity in the corresponding cell clusters from snRNA-seq and snATAC-seq respectively[28] (Fig. 1c). Furthermore, the distribution of different cell types profiled by the two methods is highly concordant across the samples, ranging from 2.5% RGC to 55.2% Rod (Fig. 1d, Supplementary Table 2).

A total of 430,567 open chromatin regions (OCRs) were identified from the snATAC-seq data, ranging from 48,764 to 199,666 per cell type (Methods, Supplementary Table 3). To assess the quality of these OCRs, we compared them with the ones from previously published bulk ATAC-seq data[29]. The snATAC-seq OCRs showed high sensitivity, capturing most OCRs identified by bulk ATAC-seq and the cell type specific OCRs that are largely missing by bulk ATAC-seq (Supplementary Fig. 1a,b,c). Specifically, 74.9% and 84.2% of OCRs identified by bulk ATAC-seq on the retina and macula tissues were detected in the snATAC-seq dataset respectively [29], and 96.2% of putative active enhancers previously identified were found in the snATAC-seq OCR list[29] (Supplementary Fig. 1a). Consistent with that Rod is the most abundant cell type in the retina, OCRs in Rod show the highest correlation with the bulk retina data with a Pearson correlation of 0.69 (Supplementary Fig. 1b). Lower correlations are observed in other cell types, particularly rare cell types, for example, a Pearson correlation of 0.41 for RGC (Supplementary Fig. 1b). Conversely, 74.0% of the OCRs are only detected by snATAC-seq, indicating a large portion of OCRs are present in a subset of cell types. Indeed, 51.5% of the snATAC-seq OCRs are unique to one cell type (Supplementary Table 2). As expected, the cell type specific OCRs are largely missed by the bulk ATAC-seq with a low detection rate of 14.3% (Supplementary Fig. 1c). To further evaluate the snATAC-seq OCRs, we examined TF binding motif enrichment in the OCRs for each cell type (Fig. 1e). Consistently, many TFs identified are previously shown to play cell type specific role in the retina, such as *OTX2, CRX, MEF2D* in photoreceptor cells, *ONECUT2* in HC, *NFIA, NFIB, NFIX, LHX2* in MG, supporting the quality of this dataset[30–34].

Putative linked cis regulatory elements (LCREs) among the OCRs were identified by calculating the correlation between the accessibility of OCRs and the nearby (+/-250kb) promoter/gene expression (Fig. 1f). As a result, about 16.6% (71,274) of the OCRs are linked to 13,405 target genes, averaging 5.9 LCREs per gene per cell type. As expected, LCREs are enriched for the CREs identified in previous studies, with 74.2% and 87.0% of the LCREs found in the ENCODE cCRE registry[6] and recent cCREs atlas[17] respectively (1.44- and 1.26-fold enrichment compared to all the OCRs, two-sided binomial test, *p* < 2.2 × 10^−16^). Furthermore, LCREs are highly enriched with active enhancers. For example, 83.8% of LCREs in Rod carry the epigenetic modifications of active enhancers, concurrent H3K4me2 and H3K27ac, a 2.1-fold enrichment compared to all the OCRs (two-sided binomial test, *p* < 2.2 × 10^−16^ Fig. 1g).

Interestingly, LCREs are depleted from cell type specific OCRs. For each cell type, on average 5.9% of LCREs are in cell type specific OCRs, 62.1% of LCREs are from OCRs shared by multiple cell types, and 32.0% of LCREs are from constant OCRs (Fig. 1g). Furthermore, LCREs tend to be in more dynamic OCRs with overall 57.3% in the differential accessible regions (DARs), a 2.2-fold enrichment compared to all the OCRs (*p* < 2.2 × 10^−16^, Supplementary Fig. 1d).

### Significant proportion of sc-eQTLs are cell type specific in retina

To profile genetic variation in the donors, WGS was performed for each donor and a total of 9.8 million genetic variants were identified after QC (Supplementary Fig. 2a). To identify genetic variants that affect gene expression, we mapped sc-eQTLs for each major retinal cell type. Due to the limited number of individuals available for our study, only variants with allele frequency ≥0.1 that are within OCRs surrounding the genes (-/+250kb of gene transcription start site, TSS) were tested, totaling 421,004 variants, averaging 59.9 variants per gene and 2.8 variants per OCR per cell type.

14,377 sc-eQTLs that reach gene level significance with false discovery rate (FDR) < 10% were identified. The variants that are in linkage disequilibrium (LD) (*r*^2^ > 0.5) from the same sc-eGene were grouped, resulting in a total of 5,688 independent sc-eQTL sets associated with 4,069 sc-eGenes, ranging from 704 to 1,175 sc-eQTL sets per cell type (Fig. 2a, b, Supplementary Table 4). The majority (86.1%-91.8%) of sc-eGenes has one sc-eQTL set per cell type (Supplementary Fig. 2b). Interestingly, most of sc-eQTLs are cell type specific, with 87.0%-92.3% identified in only one cell type (Fig. 2a). Furthermore, the remaining sc-eQTLs that are observed in multiple cell types are often shared among closely related cell types, such as between rod and cone photoreceptors (Supplementary Fig. 2c). Consistently, the effect of sc-eQTLs is correlated with the cell type similarity (Fig. 2c); for example, a stronger correlation is observed between rod and cone photoreceptors (Pearson correlation *r* = 0.6). These results suggest that the same genetic variant has a more concordant effect on gene regulation among closely related cell types, as they share a similar transcription program. Interestingly, the effect size of sc-eQTLs shared by multiple cell types in distal OCRs (which are non-promoter OCRs) is greater than that of the ones unique to one cell type (*e*.*g*., Rod, two-sided Wilcoxon rank sum test, *p* = 1.88 × 10^−5^, Fig. 2d). Consistently, sc-eQTLs shared by multiple cell types in distal OCRs are closer to gene TSS than those unique to one cell type (*e*.*g*., Rod, two-sided Wilcoxon rank sum test, *p* = 9.75 × 10^−11^, Fig. 2e).

**Fig. 2:**
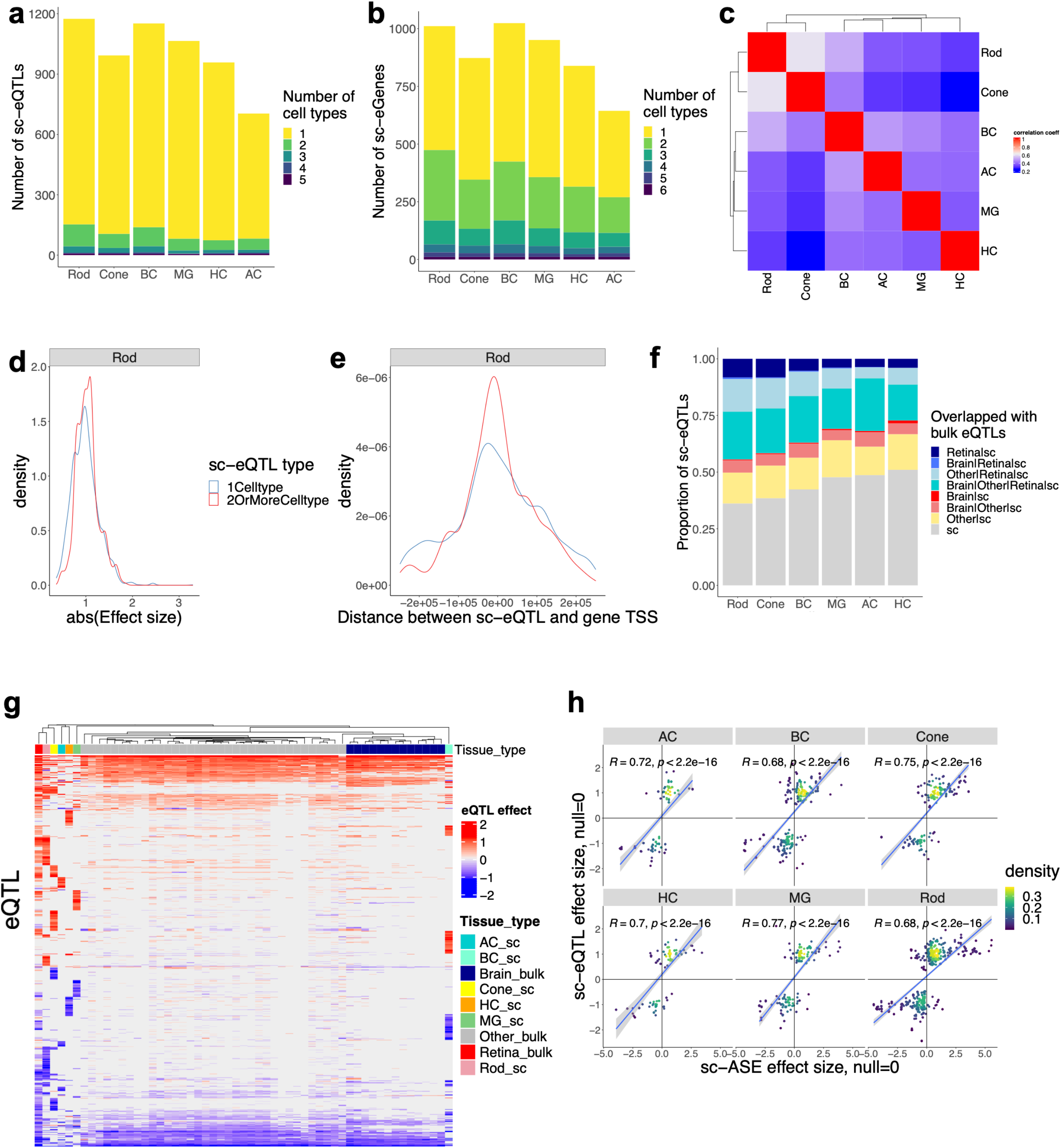
Identification of sc-eQTLs in retinal cell types. **a** The number of independent index sc-eQTLs reaching gene-level FDR < 0.1 per cell type. **b** The number of sc-eGenes reaching gene-level FDR < 0.1 per cell type. **c** Heatmap shows the Pearson correlation of sc-eQTL effect size across retinal cell types. **d** The sc-eQTLs identified in two or more cell types in distal OCRs have greater effect size than the ones identified in one cell type in distal OCRs in Rod. Two-sided Wilcoxon rank sum test, *p* = 1.88 × 10^−5^. **e** The sc-eQTLs identified in two or more cell types in distal OCRs are closer to gene TSS than the ones unique to one cell type in distal OCRs in Rod. Two-sided Wilcoxon rank sum test, *p* = 9.75 × 10^−11^. **f** The proportions of gene-level significant sc-eQTLs overlapping with gene-level significant bulk eQTLs. sc: the identified sc-eQTLs. Other: other tissue bulk eQTLs. **g** Heatmap shows effects of the sc-eQTLs and the overlapped bulk eQTLs are largely consistent across different retinal cell types and tissues. **h** The effect size of the overlapped sc-eQTLs and sc-ASEs in the corresponding cell type are significantly positively correlated. The Pearson correlation coefficient and p-values are indicated in the figure.

### Validation of sc-eQTLs with bulk eQTLs and sc-ASE

To evaluate the quality of sc-eQTLs, we compared them with the eQTLs previously identified in bulk retina and other tissues from the GTEx project[35]. sc-eQTLs are enriched for bulk eQTLs. On average 35.6% of sc-eQTLs are overlapped with the bulk retina eQTLs (4.4-fold enrichment compared to background variants, two-sided binomial test *p* < 1.2 × 10^−166^) and 56.0% overlapped with the bulk eQTLs from all the 49 tissues (2.3-fold enrichment compared to background, two-sided binomial test *p* < 2.1 × 10^−145^,, Fig. 2f). The proportion of overlap varies among cell types (Fig. 2f). As expected, the highest overlap (63.9%) is observed for the most abundant cell type, Rod, while the lowest overlap is observed for HC at 49.0% (Fig. 2f). Effect direction of eQTLs across different cell types and tissues is largely concordant (Fig. 2g).

We further validated these sc-eQTLs with sc-ASEs. sc-eQTLs are enriched for sc-ASEs. sc-ASEs are detected in 18.8%-34.0% of the sc-eQTLs that were tested for sc-ASEs (with the highest overlapping in Rod, 34.0%), on average 2.5-fold enrichment compared to background variants (two-sided binomial test *p* < 1.2 × 10^−12^, Supplementary Fig. 2d). The effect size and direction are positively correlated (Pearson correlation, *r* in 0.68-0.77, *p* < 2.2 × 10^−16^), with the majority (82.5%-94.2%) of the overlapped variants having the same direction (Figure 2h, Supplementary Fig. 2d). Altogether, these results support that the majority of sc-eQTLs identified are likely to be true positives.

### Cell type specific sc-eQTLs often reside in OCRs shared by multiple cell types

An interesting observation is that most (87.0%-92.3%) of sc-eQTLs are unique to one cell type, while the associated sc-eGenes (94.6%-98.9%) are almost always expressed in multiple cell types (Fig. 2a, 3a). Specifically, only a small proportion (1.8%-6.0%) of sc-eQTLs and their associated sc-eGenes share the same pattern of cell type specificity. In over 90% of the cases, while the sc-eQTL is observed in one or a subset of cell types, the sc-eGenes are expressed in multiple cell types. Interestingly, for the same sc-eGene, different sc-eQTLs are often observed in different cell types (36.4% of total sc-eQTLs) (Fig. 3b), and these sc-eQTLs tend to be in different OCRs (34.0% of total sc-eQTLs, Supplementary Fig. 2e). This does not result from cell type specific accessibility of the OCRs, as OCRs are often accessible in multiple cell types while sc-eQTL effect of the resident variants are only observed in one or subset of cell types. This is not due to the differential accessibility of the OCRs as well, since only a small proportion (8.8%-19.4%) of sc-eQTLs in the DARs of the corresponding cell types. Only a small fraction (11.4%) of sc-eQTLs reside in OCRs whose accessibility have matching cell type specificity as those of the sc-eQTLs (Fig. 3c). For example, the variant rs10793810 is a MG specific sc-eQTL of SLC27A6, and enhances the binding of FOXP2 (highly expressed in MG), to a MG-specific enhancer of SLC27A6 (Fig. 3d). In contrast, most (89.1%) of sc-eQTLs are within the OCRs shared among multiple cell types (Fig. 3c), suggesting that modulation of gene expression by genetic variants is primarily driven by activity of trans-factors such as cell type specific TFs, rather than the accessibility of cis-elements. For example, the variant rs62308155 is identified as a Rod specific sc-eQTL of REST, likely through disrupting the binding of NR3C1, which is highly expressed in Rod but minimally in Cone, to an enhancer accessible in both Rod and Cone (Fig. 3e). Supporting the roles of trans-factors in driving cell type specific sc-eQTL effect genome-widely, the TFs, whose motifs are perturbed by genetic variants, have higher expression in the cell types where the variants have sc-eQTL effect, compared to the cell types where the variants do not have effect (*e*.*g*., Rod, one-sided Wilcoxon rank sum test, *p* < 4.7 × 10^−6^, Fig. 3f).

**Fig. 3:**
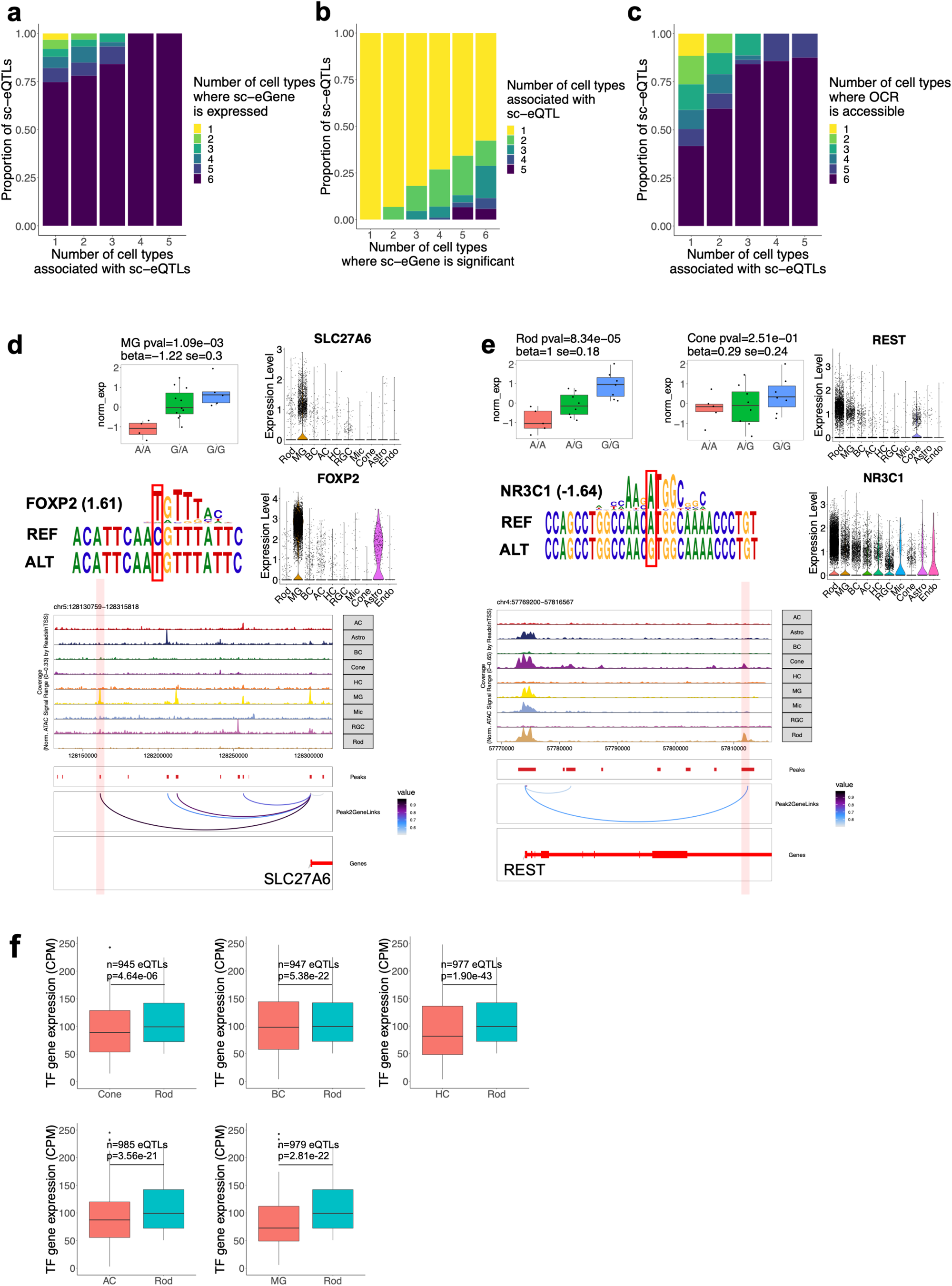
Cell type specific sc-eQTLs are often within OCRs shared by multiple cell types. **a** The sc-eGenes of sc-eQTLs are often expressed in multiple cell types. **b** sc-eQTLs of the same sc-eGene are often different across cell types. **c** The majority of the sc-eQTL-containing OCRs are accessible in multiple cell types. **d** The variant rs10793810 is a MG-specific sc-eQTL of SLC27A6 and located in a MG-specific OCR. This variant is predicted to enhance the binding of FOXP2, which is highly expressed in MG. This OCR is a predicted LCRE of SLC27A6. **e** The variant rs62308155 is a Rod-specific sc-eQTL of REST, and resides in an OCR accessible in Rod and Cone. This variant is predicted to disrupt the binding of NR3C1, which is highly expressed in Rod but minimally in Cone. This OCR is a predicted LCRE of REST. **f** The TFs, whose binding sites are perturbed by a variant that is sc-eQTL in Rod but not in another cell type, have higher expression in Rod than the other cell type. One-sided Wilcoxon rank sum test. The p-value and sample size n are indicated in the figure. The Y axis was set between 0 and 250 for better visualization of the data.

### Significant proportion of sc-caQTLs are cell type specific in retina

In parallel with sc-eQTL analysis, to identify genetic variants that affect chromatin accessibility, we performed sc-caQTL analysis by examining the association between each OCR and the common variants within it for each major retinal cell type. A total of 174,419 OCRs (ranging from 54,716 to 95,020 OCRs per cell type) and the same set of variants tested for sc-eQTLs were analyzed (Methods). Upon genome-wide multiple testing corrections, a total of 23,287 sc-caQTLs were identified (FDR < 10%), which were grouped into 12,482 independent sc-caQTLs sets mapped in 10,298 OCRs based on LD (*r*^2^ > 0.5), ranging from 391 to 4,789 sc-caQTLs sets per cell type (Fig. 4a,b and Supplementary Table 5). The majority (88.0%) of sc-caQTL-containing OCRs, namely sc-caQTL-associated peaks (sc-caPeaks) in this study, contain only one sc-caQTL set (Supplementary Fig. 3a). The majority of sc-caQTLs are cell type specific with 62.3%-85.7% unique to one cell type, a lesser degree compared to sc-eQTLs. Similar to sc-eQTLs, the effect sizes of sc-caQTLs are correlated across cell types, with stronger correlation observed between more closely related cell types (Fig. 4c and Supplementary Fig. 3b). The distal sc-caQTLs common in multiple cell types have significantly greater effect than the ones unique to one cell type (*e*.*g*., Rod, one-sided Wilcoxon rank sum test, *p* < 2.2 × 10^−16^, Fig. 4d).

**Fig. 4:**
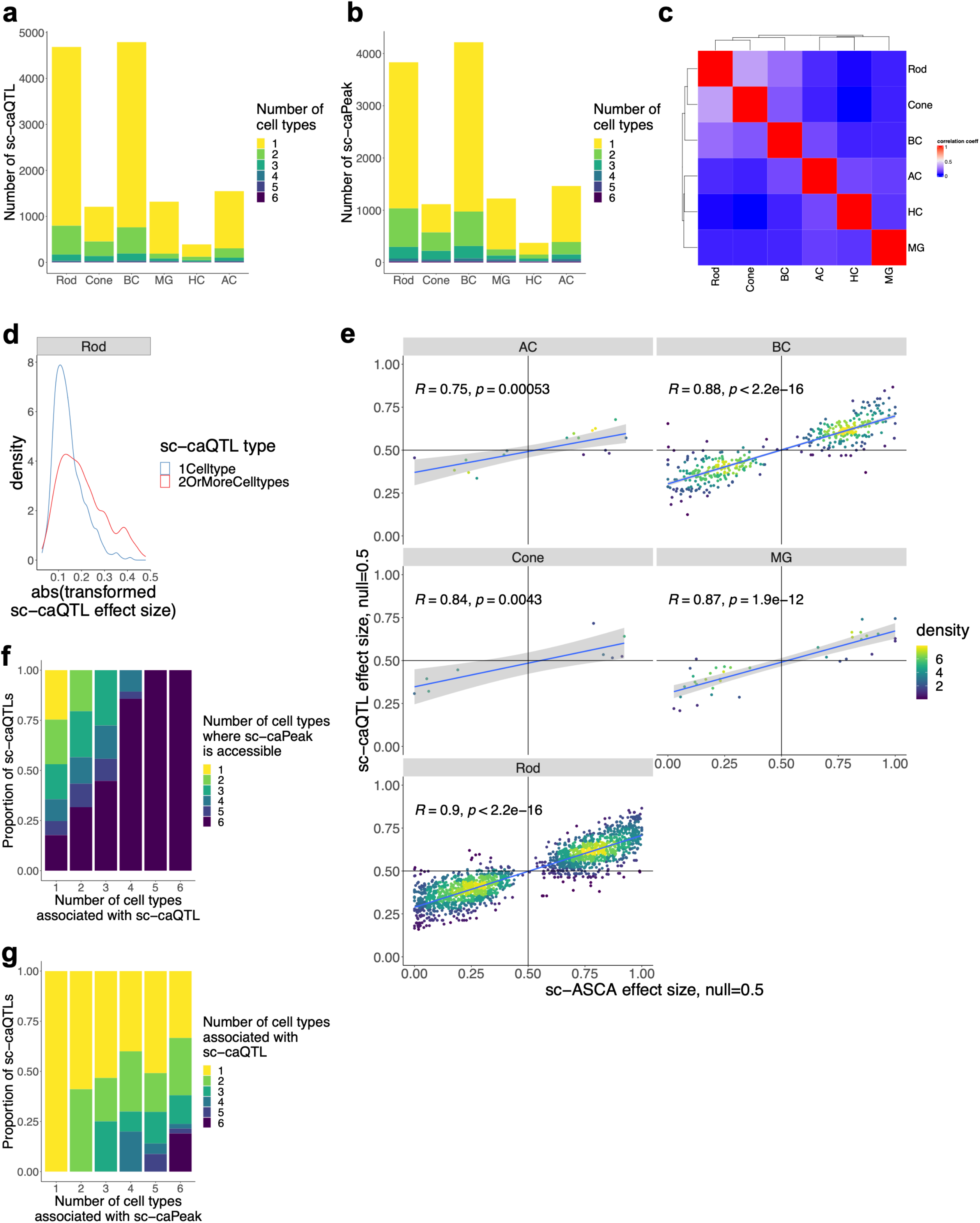
Identification of sc-caQTL in retinal cell types. **a** The number of independent index sc-caQTLs reaching genome-level FDR < 0.1 per cell type. **b** The number of sc-caPeak reaching genome-level FDR < 0.1 per cell type. **c** Heatmap shows the Pearson correlation of sc-caQTL effect size across retinal cell types. **d** The distal sc-caQTLs identified in two or more cell types have greater effect size than the ones identified in one cell type in Rod. Two-sided Wilcoxon rank sum test, *p* < 2.2 × 10^−16^. **e** The effect size of sc-ASCA and the population effect size of sc-caQTLs is significantly positively correlated for each cell type. The Pearson correlation coefficient and p-values are indicated in the figure. **f** The majority of the sc-caPeaks (i.e. sc-caQTL-containing OCRs) are accessible in multiple cell types. **g** The sc-caQTLs of the same sc-caPeak can be different across cell types.

### Validation of sc-caQTLs with sc-ASCA

To assess the quality of the sc-caQTLs identified, we compared sc-caQTLs with sc-ASCAs. sc-ASCAs are detected in 8.7%-41.8% of the sc-caQTLs that were tested for sc-ASCAs (with the highest overlapping rate in Rod, 41.8%), on average 15.9-fold enrichment compared to background (two-sided binomial test, *p* < 6.8 × 10^−11^, Supplementary Fig. 3c). Furthermore, the effect size and direction of sc-ASCAs and the overlapped sc-caQTLs are positively correlated (Pearson correlation, *r* in 0.75-0.90, *p* < 5.4 × 10^−4^), with the majority (82.4%-100%) of the overlapped variants having the same direction (Fig. 4e, Supplementary Fig. 3c). These results support that the sc-caQTLs identified are indeed enriched of variants associated with change in chromatin accessibility. Conversely, 33.3%-54.5% of the identified sc-ASCAs overlap with sc-caQTLs depending on cell type. Interestingly, the size of OCRs containing sc-ASCA-only variants (not overlapped with sc-caQTL) are significantly larger than the ones containing variants which are both sc-ASCA and sc-caQTL (Supplementary Fig. 3d, *e*.*g*., one-sided wilcoxon rank sum test, *p* < 1.47 × 10^−21^ in Rod). This observation suggests that the variants in wider OCRs tend to have local effect, while the variants in the narrow OCRs are more likely to affect accessibility of the entire OCRs.

### Cell type specific sc-caQTLs can reside in OCRs accessible in multiple cell types

Like sc-eQTLs, most (62.3%-85.7%) of sc-caQTLs are unique to one cell type, while the majority (74.8%) of sc-caPeaks are accessible in multiple cell types (Fig. 4a, f). Specifically, 24.4% of sc-caQTLs and their caPeaks share the same pattern of cell type specificity. 75.6% of sc-caQTLs are found in one or a subset of cell types while the sc-caPeaks are accessible in multiple cell types (Fig. 4f). Furthermore, the cell type unique sc-caQTLs is not due to the differential accessibility of OCRs alone, since only a small proportion (14.9%-34.3%) of sc-caQTLs were observed in the DARs of the corresponding cell types. Interestingly, for the sc-caPeaks common in multiple cell types, different sc-caQTLs variants are observed in different cell types, accounting for 13.9% of total sc-caQTLs (Fig. 4g). As an example where the cell specificity matches between sc-caQTLs and their residing OCRs, the variant rs12447029 has MG specific sc-caQTL effect through strengthening the binding of NFE2L2, which is highly expressed in multiple cell types, to a MG-specific enhancer (Fig. 5a). Consistently, the corresponding OCR is a LCRE of GRIN2A, and rs12447029 is a sc-eQTL for GRIN2A in MG (Supplementary Fig. 3e). In contrast, the cell type specificity of the vast majority of sc-caQTLs cannot be explained by the cell type specificity of the corresponding OCRs alone. 68.3% of the sc-caQTLs are unique to one cell type but reside in the OCRs observed in multiple cell types, indicating the modulation of chromatin accessibility by genetic variants is often cell type-dependent, probably through affecting the binding of cell type specific trans-factors (Fig. 4f). For example, although accessible in Rod, Cone and BC, the variant rs6859300 affects the chromatin in Rod only, possibly through enhancing the binding of EPAS1, which is highly expressed in Rod while lowly expressed in Cone and BC (Fig 5b). Consistently, the corresponding OCR is a LCRE of WWC1, and rs6859300 is a sc-eQTL of WWC1 in Rod (Supplementary Fig. 3f). Furthermore, the TFs, whose motifs are perturbed by genetic variants, have higher expression in the cell types where the variants have sc-caQTL effect, compared to the cell types where the variants do not have effect, supporting the role of trans-factors in driving cell type specific sc-caQTL effect genome-widely (*e*.*g*., Rod, one-sided Wilcoxon rank sum test, *p* < 4.5 × 10^−19^, Fig. 5c).

**Fig 5:**
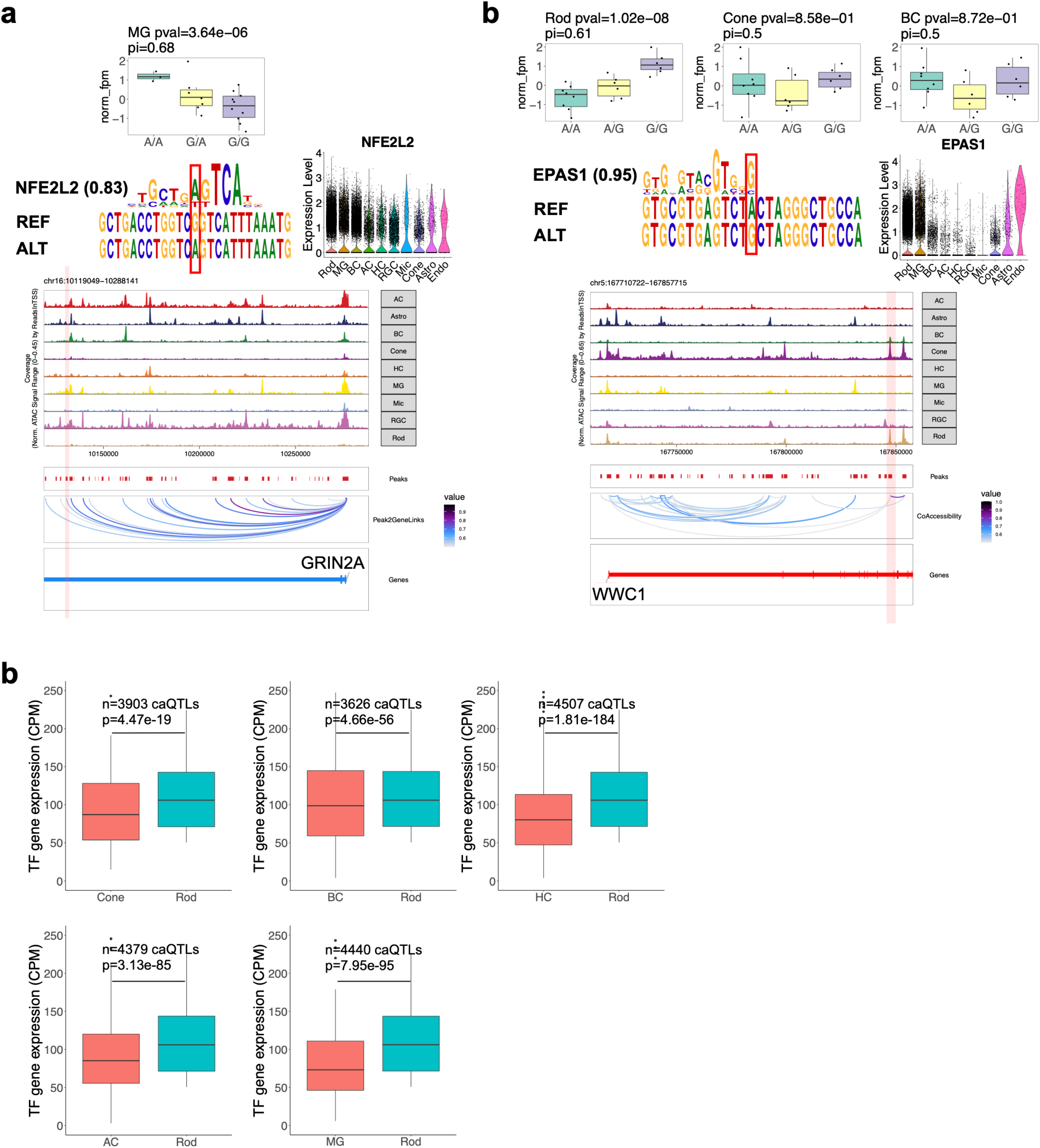
Examples of cell type specific sc-caQTLs. **a** The variant rs12447029 is a MG-specific sc-caQTL of its residing OCR, and resides in a MG-specific OCR. This variant is predicted to enhance the binding of NFE2L2, increasing chromatin accessibility of its residing OCR in MG. NFE2L2 is highly expressed in multiple cell types. This OCR is a predicted LCRE of GRIN2A. **b** The variant rs6859300 is a Rod-specific sc-caQTL of its residing OCR, and resides in an OCR accessible in Rod, Cone, and BC. This variant is predicted to enhance binding of EPAS1, increasing chromatin accessibility of its residing OCR in Rod. EPAS1 is highly expressed in Rod but lowly in Cone and BC. This OCR is a predicted LCRE of WWC1. **c** The TFs, whose binding sites are perturbed by a variant that is sc-caQTL in Rod but not in another cell type, have higher expression in Rod than the other cell type. One-sided Wilcoxon rank sum test. The p-value and sample size n are indicated in the figure. The Y axis was set between 0 and 250 for better visualization of the data.

### Interaction among OCRs

Previous studies suggest that multiple regulatory elements can be regulated by a single genetic variant[12]. One possible mechanism is that the accessibility of a “master” element affects the accessibility of neighboring “dependent” elements[12]. To examine this phenomenon in our dataset, we identified 2511 dependent regions associated with 1942 master regions (Methods). Among them, 360 master regions that are LCREs, are associated with 427 dependent regions that are LCREs of the same genes. The proportions of sc-caQTLs associated with the dependent OCRs (*e*.*g*., 1.8-fold enrichment compared to background variants in Rod, two-sided binomial test, *p* = 1.48 × 10^−88^) and dependent LCREs (*e*.*g*., 1.7-fold enrichment compared to background in Rod, two-sided binomial test, *p* = 4.49 × 10^−12^) are significantly enriched compared to background variants respectively, suggesting the association between sc-caQTL/master elements and dependent elements are not random (Fig. 6a). The effect size of sc-caQTLs on the master regions and dependent regions are positively correlated (an average correlation coefficient of 0.60, *p* = 1.5 × 10^−7^), with the majority (65.0%-82.0%) of the sc-caQTLs having the same effect direction on the master and dependent regions (Fig. 6b). Furthermore, slightly higher enrichment in DARs and active enhancer modifications (the concurrent H3K27ac and H3K4me2) was observed in the master regions than the dependent regions (Fig. 6c).

**Fig. 6:**
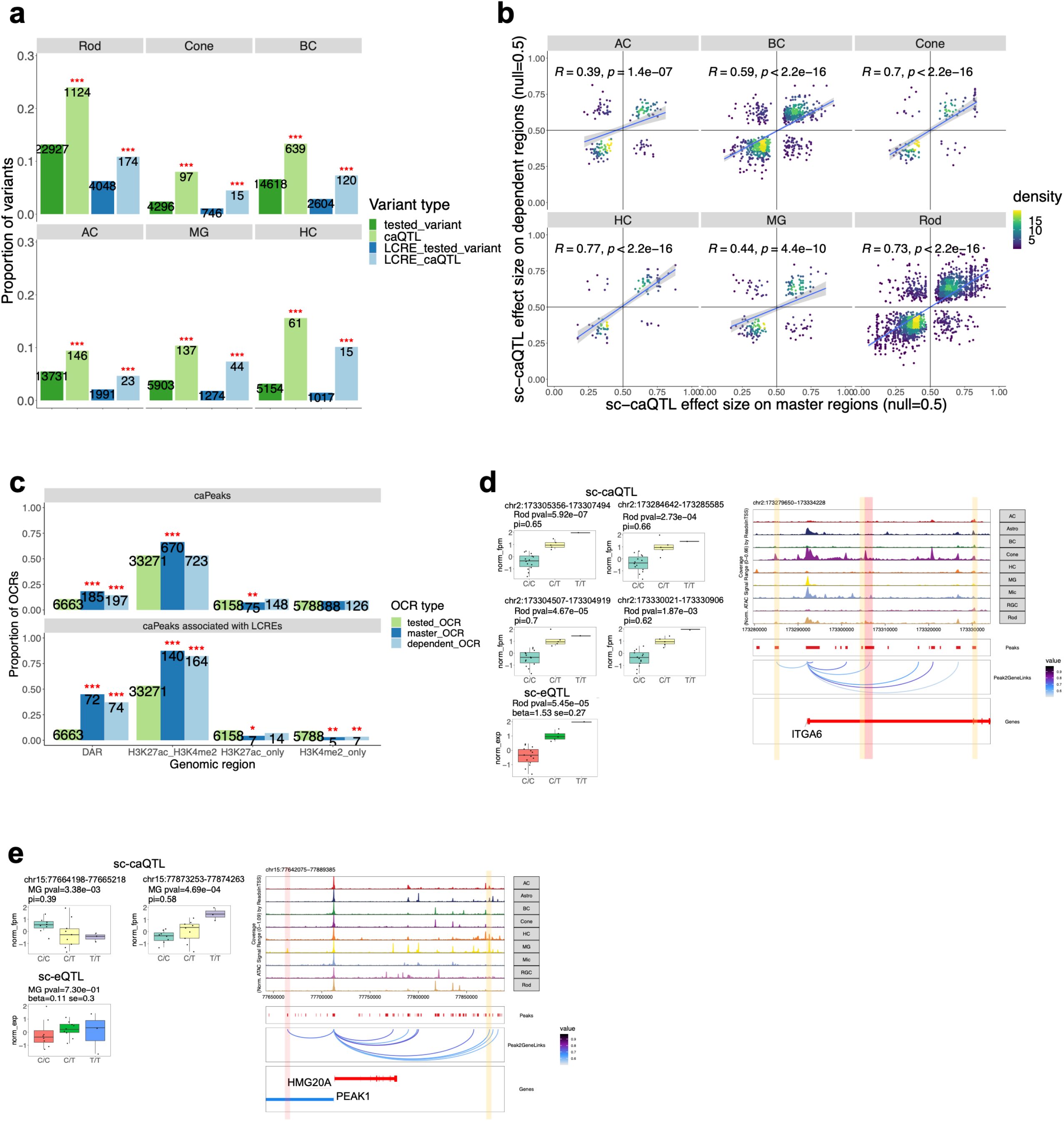
The sc-caQTLs with effects on multiple genomic regions. **a** The proportion of variants affecting dependent OCRs and the proportion of variants affecting dependent LCREs. The proportions associated with sc-caQTLs are significantly higher than the proportions associated with background variants. The numbers of variants affecting dependent OCRs are indicated in the figure. **b** The effect size of sc-caQTLs on master regions and the effect size of sc-caQTLs on dependent regions are positively correlated. **c** The proportion of OCRs that are DARs and have concurrent H3K27ac and H3K4me2 modifications in Rod. The numbers of caPeaks with different features are indicated in the figure. **d** The variant rs7596259 is a sc-caQTL and affects a master region (red) and three dependent regions (yellow) in the same effect direction in Rod. The master region and one dependent region are LCREs of ITGA6. This variant is also a sc-eQTL of ITGA6 in Rod. **e** The variant rs1493699 is a sc-caQTL and affects a master region (red) and a dependent region (yellow) in the opposite direction in MG. Although the two regions are LCREs of PEAK1, this variant is not a sc-eQTL of PEAK1 in MG.

Although the majority (66.5%-87.7%) of the master regions have one dependent region, some have multiple dependent regions. For example, the sc-caQTL variant rs7596259 increases accessibility of its residing master region, and is associated with the increased accessibility of the other three dependent regions in Rod (Fig. 6d). This sc-caQTL is also a sc-eQTL and increases the gene expression of ITGA6 in Rod, suggesting some of the affected regions might be important for gene expression regulation (Fig. 6d). Indeed, the master region (chr2:173305356-173307494) and one dependent region (chr2:173284642-173285585) are the predicted LCREs of ITGA6 (Fig. 6d). Moreover, the sc-caQTLs affecting multiple regions in the same effect direction are more likely to overlap with sc-eQTLs in the corresponding cell type than the sc-caQTLs affecting multiple regions in different effect directions (in Rod 15.9% vs. 4.2%, two-sided binomial test *p* = 2.25 × 10^−8^), which might be due to compensation between the elements with opposite effect directions canceling out their impact on gene expression. For example, the sc-caQTL variant rs1493699 reduces the accessibility of its residing master region (chr15:77664198-77665218), and is associated with the increased accessibility of a dependent region (chr15:77873253-77874263) in MG (Fig. 6e). Although the two elements are LCREs of PEAK1, this sc-caQTL is not a sc-eQTL of PEAK1, suggesting that these regulatory elements might compensate for each other and overall do not change gene expression (Fig. 6e).

### Prioritizing causal variants and cell type context underlying GWAS loci

The single cell multiomics dataset provides opportunities to fine map GWAS loci in a cell type context. We first investigated the cell type enrichment of GWAS loci associated with 11 eye traits or disorders[36–42] based on cell type chromatin accessibility and gene expression respectively[43][44–47] (Methods). Interestingly, the GWAS loci enrichment identified from chromatin state and gene expression converges to the same cell types (Fig. 7a,b, and Supplementary Fig. 4a, Benjamini-Hochberg correction, *p. adj* < 0.1). Specifically, primary open-angle glaucoma (POAG) related traits, such as cup areas (CA) and vertical cup-disc ratio (VCDR) of optic nerve, intraocular pressure (IOP), and POAG, displayed enrichment in the DARs, OCRs, and/or genes expressed in astrocytes and MG (*p* < 9.7 × 10^−3^, *p. adj* < 0.1, Fig. 7a,b). Refractive error and myopia loci[42], displayed enrichment in the DARs, OCRs, and/or genes expressed in most of major retinal cell types (*p* < 8.2 × 10^−3^, *p. adj* < 0.1) (Fig. 7a,b and Supplementary Fig. 4a). The loci associated with choroid/retina disorders, retinal detachments/breaks, and retinal problems[41], showed enrichment in the DARs of MG (Fig. 7a, *p* < 7.2 × 10^−3^, *p. adj* < 0.1).

**Fig. 7:**
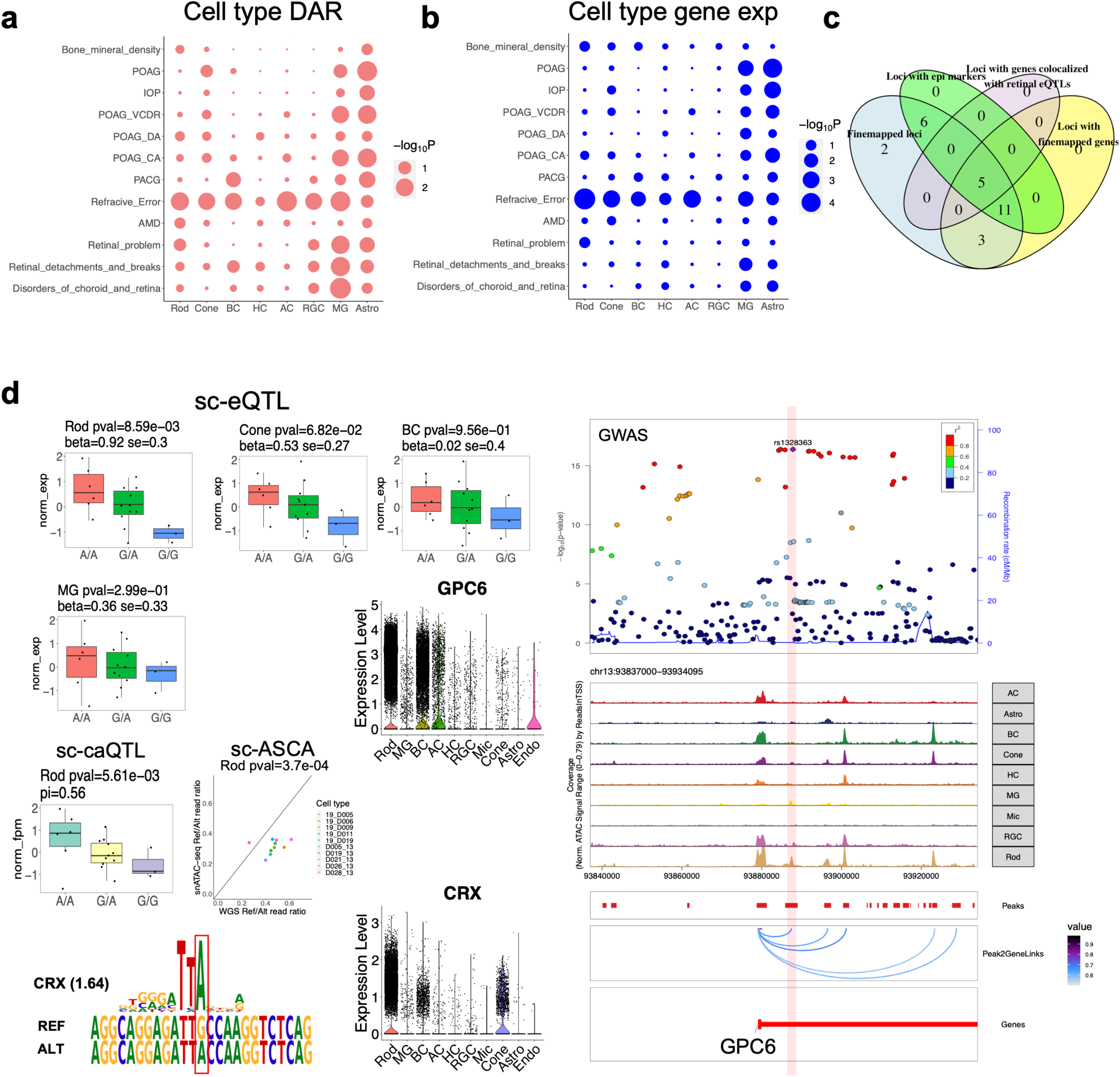
Cell type enrichment and causal variant prioritization underlying GWAS loci. **a** The cell type enrichment of 11 eye-related and one control GWAS loci by partitioning the heritability enrichment in cell type DARs with LDSC. POAG: primary open-angle glaucoma. IOP: intraocular pressure. VCDR: vertical cup-disc ratio of optic nerve. CA: cup area of optic nerve. DA: disc area of optic nerve. PCAG: primary angle closure glaucoma. AMD: age-related macular degeneration. **b** The cell type enrichment of 11 eye-related and one control GWAS loci based on gene expression cell type specificity from snRNA-seq data with MAGMA.Celltyping. **c** Venn diagram showing the features of the prioritized GWAS loci overlapped with sc-QTL and/or sc-ASCA. **d** The variant rs1328363 associated with refraction error and myopia with PIP=0.308 is a sc-eQTL of GPC6, a sc-caQTL of its residing OCR and a sc-ASCA in Rod. Its residing OCR is accessible in Rod, Cone, BC and MG, and a predicted LCRE of GPC6. This variant strengthens the binding of a photoreceptor-specific TF (CRX). This variant is also a marginal sc-eQTL of GPC6 in Cone, consistent with CRX also being a TF for Cone, much lower expression of GPC6 and lower accessibility of the corresponding LCRE in Cone.

To identify causal variants and target genes with a cellular context underlying GWAS loci, we fine-mapped GWAS variants associated with three eye diseases, glaucoma[36], age-related macular degeneration[40], and refraction error/myopia[42]. We incorporated functional annotation (including OCR and LCRE derived from single cell multiomics data) of variants to prioritize GWAS loci[48]^56^. As a result, 818 variants with posterior inclusion probability (PIP) > 0.1 were identified, which contain potential causal variants and are enriched of variants in regulatory regions (Supplementary Fig. 4b,c,d). Among them, 27 variants are sc-caQTL, sc-eQTL, and/or sc-ASCA (Fig. 7c, Supplementary Fig. 4e,f, Supplementary Table 6). 22 (81.5%) of these 27 variants are in the regions with epigenetic modifications, H3K27ac and/or H3K4me2, supporting their regulatory role (Supplementary Table 6). To identify the target genes, 19 of the 27 variants were linked to 24 target genes through sc-eQTLs, LCREs and gene annotation (Supplementary Table 7). As expected, 14 (58.3%) of the 24 candidate genes are the nearest genes adjacent to the variants. Furthermore, 6 of these 24 genes are also supported by the colocalization of GWAS signals with retinal bulk eQTL signals. For example, the variant rs511217 is a fine-mapped variant associated with refraction error and myopia (PIP= 0.176). This variant is a sc-eQTL of KCNA4 and a nominal significant sc-caQTL of its residing OCR in BC. The corresponding OCR is a predicted LCRE of KCNA4. Consistently, the GWAS signal is colocalized with the retinal bulk eQTL signal of KCNA4 as well (Supplementary Fig. 5).

Our integrative analysis also provided potential insights for the cell type specific regulatory mechanisms of GWAS loci (Fig 7d). For example, rs1328363 is a fine-mapped variant associated with refraction error and myopia (PIP=0.308). This variant may achieve Rod specific effect (a sc-ASCA, nominal significant sc-eQTL, and nominal significant sc-caQTL) in increasing expression of GPC6 through strengthening the binding of a photoreceptor-specific TF (CRX) to a GPC6 enhancer which is accessible in multiple cell types (Fig 7d). This variant is also a marginal sc-eQTL in Cone, concordant with CRX also being a TF for Cone, much lower expression of GPC6, and lower accessibility of the corresponding enhancer in Cone. GPC6 encodes a putative cell surface glypican coreceptor, implicating its role in controlling cell growth and division.

## Discussion

In this study, by combining single-cell multiomics to profile cells from human retina and genomic sequencing, we identified regulatory elements, mapped effect of genetic variants, and elucidated modulation mechanisms underlying gene regulation in individual cell type contexts *in vivo*. The genetic effects on gene expression measured by sc-eQTLs and sc-ASE are highly concordant, while the gene effects on chromatin accessibility assessed by sc-caQTLs and sc-ASCAs also show consistency. Additionally, sc-eQTLs are enriched of bulk eQTLs from retina and other tissue types, and higher overlapping rate was observed for the sc-eQTLs identified in the most abundant cell type or the ones common in multiple cell types. Altogether, these results support the quality of the mapped genetic effects on gene expression and chromatin accessibility. Interestingly, a significant proportion (44.0%) of sc-eQTLs are missed from bulk eQTLs, which might be due to most of the sc-eQTLs being cell type specific, thus the cell type specific signals, in particular the ones associated with rare cell types, might be diluted and not detectable in the bulk level. It is also likely that some sc-eQTLs have opposite effect direction in different cell types, so the overall effect in the bulk level is canceled out, although we observed a very small proportion of sc-eQTLs in such cases.

Intriguingly, most of the mapped sc-eQTLs and sc-caQTLs are cell type specific, while most of eGene and caPeaks are active in multiple cell types, suggesting genetic variants modulate gene expression and chromatin state in a cell type dependent manner. Surprisingly, the majority of cell type specific sc-eQTLs and sc-caQTLs reside in the regulatory elements accessible in multiple cell types. Furthermore, the TFs, whose motifs are perturbed by genetic variants, have higher expression level in the cell types where the variants have cell type specific effect, compared to the cell types where the variants do not have effect. Altogether, our study suggested that for the cell types sharing a similar lineage, cell type specificity of genetic effect is not primarily due to cell type specificity of the affected cis-elements, but may be mainly achieved by perturbing the binding of cell type specific trans-factors (Fig. 8). Specifically, we hypothesized that some regulatory genomic regions in the cells sharing a similar lineage may be first opened and primed by pioneer factors, thus different cell types could have common OCRs, and these OCRs can be bound by additional different trans-factors later depending on cell type/state context, in a collaborative manner. Therefore, genetic variants affecting the binding of cell type/state specific trans-factors within the common OCRs could have cell type specific effect on gene expression and chromatin accessibility. These results also suggested the accessibility of a genomic region does not necessarily indicate its activity, and an accessible regulatory element may be inactive and could be activated by the binding of additional trans-factors in a given cell type/state context. However, for the cell types from different lineages, the affected cis-elements may play important role in determining the cell type specificity of genetic variant effects, which needs further investigation.

**Fig. 8:**
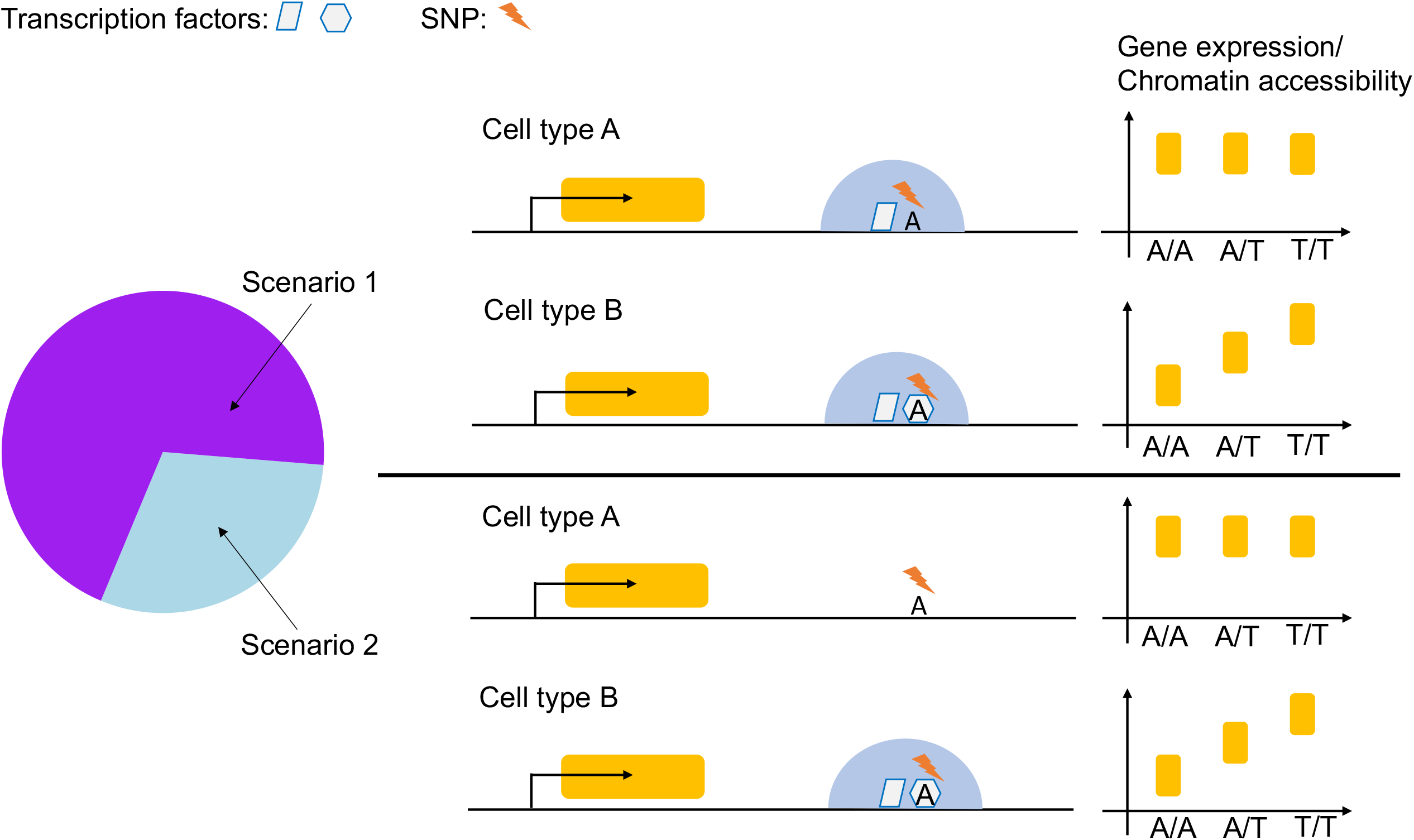
A model of the cell type specific effect of genetic variants. The schematic plot shows that for the cell types sharing a similar lineage, the cell type specific effect of genetic variants is not primarily due to cell type specific chromatin accessibility of cis-elements but may be mainly driven by perturbing the binding of cell type specific trans-regulators, indicating hierarchical transcription factors collaboration may play important role in cell type specific effects of genetic variants on gene regulation.

Moreover, we showed that integration of single cell multiomics and GWAS studies can increase the power to prioritize effective cell context, causal variants and genes, and better dissect the underlying regulatory mechanisms. In our study, the cell type enrichment of GWAS traits measured by gene expression and chromatin accessibility converged to the same cell types, supporting the accuracy of our result, and suggesting some GWAS loci may indeed affect regulatory elements linked to gene expression in specific cell type context. Intriguingly, our analyses showed that astrocyte and MG play important role in POAG, and MG may be involved in choroid/retina disorders, suggesting non-neuronal cell types, particularly glia cells, may be critical for neuronal diseases. MG and astrocyte are macroglia cells in the retina and play essential roles in maintaining the homeostasis and proper function of the retinal neurons[49]. In particular, astrocytes are located in the nerve fiber and ganglion cell layers, support the structure and physiology of the optic nerve head axon and modulate the extracellular matrix under elevated IOP[50], supporting their important role in glaucoma. Furthermore, we fine-mapped GWAS loci based on functional annotation of genetic variants, which prioritize the variants in regulatory regions as candidate causal variants. By overlapping the fine-mapped GWAS variants with sc-eQTL and LCREs, we identified the genes potentially contributing to myopia/refraction error and glaucoma. Moreover, combining gene expression, chromatin accessibility, and their variation driven by genetic variants in cell type context, we explained the cell type specific regulation mechanism underlying GWAS loci, which could be related to cell type specific trans-factor binding and/or cis-elements. These findings could facilitate the understanding of pathogenic mechanisms and provide guidance for functional analysis of GWAS loci and development of disease treatment.

## Conclusions

We conducted the first systematic study of how common genetic variants modulate gene expression and chromatin accessibility in major cell types of the human retina through integrative single-cell multiomics analysis. Our findings suggest effects of genetic variants on gene regulation are highly context dependent. For the cell types sharing a similar lineage, the cell type specific genetic effects may be mainly driven by trans-factors rather than the chromatin stat of the affected cis-elements. These results indicate hierarchical transcription factors collaboration may play an important role in genetic regulation of gene expression and chromatin. Our study provides novel insights on the mechanisms of gene regulation at a nucleotide level of cellular resolution, which may shed light on understanding and treating human diseases.

## Methods

### Human retina sample collection

Samples included in this study were retinal tissues of 20 donors from the Utah Lions Eye Bank (Supplement Table 1). All donors were screened for medical history, and only the ones with no records of retinal diseases were used in this study. Post-mortem phenotyping with OCT were performed to confirm that there were no drusen, atrophy, or any other disease phenotypes on retina by our previous approach[51]. One eye was collected from each donor. All eye tissues were collected and dissected within 6 hours post-mortem, according to previous protocol[52]. With 4mm and 6mm disposable biopsy punches, macula and peripheral retina were collected and flash-frozen in liquid nitrogen, and stored at -80°C before nuclei isolation. All tissues were de-identified under HIPAA Privacy Rules. Institutional approval for the consent of patients for their tissue donation was obtained from the University of Utah and conformed to the tenets of the Declaration of Helsinki.

### Nuclei isolation and sorting

Nuclei for snRNA-seq were isolated by fresh-made pre-chilled RNase-free lysis buffer (10mM Tris-HCl, 10mM NaCl, 3mM MgCl2, 0.02% NP40). The frozen tissue was resuspended and triturated in lysis buffer and homogenized with a Wheaton™ Dounce Tissue Grinder. Isolated nuclei were filtered with a 40μm Flowmi Cell Strainer. DAPI (4’,6-diamidino-2-phenylindole, 10μg/ml) was added before loading the nuclei for fluorescent cytometry sorting with a BD (Becton Dickinson, San Jose, CA) Aria II flow sorter (70μm nozzle). The sorted nuclei are ready for snRNA-seq.

Nuclei for snATAC-seq were isolated in fresh-made pre-chilled lysis buffer (10mM Tris-HCl, 10mM NaCl, 3mM MgCl2, 0.02% NP40, 1% BSA). Similar to the nuclei isolation process for snRNA-seq, frozen tissue was homogenized with a Dounce Tissue Grinder until no tissue pieces were visible. Nuclei were then washed (wash buffer: 10mM Tris-HCl, 10mM NaCl, 3mM MgCl2, 1% BSA) twice in a pre-coated (coating buffer: 10mM Tris-HCl, 10mM NaCl, 3mM MgCl2, 4% BSA) 5ml round-bottom Falcon tube (Cat. NO. 352054) at 500g, 4°C for 5min. Nuclei were resuspended in 1X diluted nuclei buffer (10X PN-2000153, PN-2000207) for a final concentration of 3000-5000 nuclei/ul.

### Single-nuclei sequencing

Single cell Gene Expression Library was prepared according to Chromium Next GEM Single Cell 3’ Reagent Kits v3.1 (10x Genomics). In Brief, single nuclei suspension, reverse transcription (RT) reagents, Gel Beads containing barcoded oligonucleotides, and oil were loaded on a Chromium controller (10x Genomics) to generate single cell GEMS (Gel Beads-In-Emulsions) where full length cDNA was synthesized and barcoded for each single cell. Subsequently the GEMS are broken and cDNA from each single cell are pooled. Following cleanup using Dynabeads MyOne Silane Beads, cDNA is amplified by PCR. The amplified product is fragmented to optimal size before end-repair, A-tailing, and adaptor ligation. Final library was generated by amplification. After quantification with KAPA Library Quantification kit (Roche), libraries were sequenced on a Novaseq 6000 Sequencer (Illumina).

Single cell ATAC Library was prepared according to Chromium Next GEM Single cell ATAC Reagent kit v1.1 (10x Genomics). In Brief, prepared nuclei were incubated with transposome and transposase entered and preferentially fragmented DNA in open region of chromatin. Transposed single nuclei, a master mix, Gel Beads containing barcoded oligonucleotides, and oil were loaded on a Chromium controller (10x Genomics) to generate GEMS (Gel Beads-In-Emulsions) where barcoded single strand DNA was synthesized. Subsequently the GEMS are broken and pooled. Following sequential cleanup using Dynabeads MyOne Silane Beads and SPRI beads, barcoded DNA fragments are amplified by PCR to generate indexed library. After quantification with KAPA Library Quantification kit (Roche), libraries were sequenced on a Novaseq 6000 Sequencer (Illumina).

### Whole genome sequencing

1 ug genomic DNA was sheared with Covaris for 70 seconds and the purification was performed with Ampure XP beads. After end repair and A-tailing, the indexed adaptors were added to the product, and subsequently purified with Ampure XP beads. The diluted library was then sequenced in an Illumina Novaseq6000 Sequencer.

### WGS data processing

The WGS variant calling followed the GATK pipeline for analyzing small sample cohorts (https://gatk.broadinstitute.org/hc/en-us/articles/360035890411-Calling-variants-on-cohorts-of-samples-using-the-HaplotypeCaller-in-GVCF-mode). Briefly, WGS data was aligned to the human reference genome (build hg19) with BWA-MEM[53]. After removing duplicate reads with MarkDuplicates (Picard) from GATK, the bam files were realigned with base quality score recalibration and local realignment with GATK4[54]. With the realigned bam files, the variants were called to generate genome-wide genotype-per-site data for each sample (gVCF). The joint genotyping was performed on variants of all samples using GATK GenotypeGVCFs. Variants from joint genotyping underwent variant recalibration with GATK. The WGS variants were then QC and filtered (Supplementary Note), and a total of 9,792,238 variants were obtained for downstream analysis.

### Quality control of sample genotypes

The sample genotypes were QC using multidimensional scaling (MDS) analysis of plink with the genotype data from the Hapmap project[55,56] (including 84 CHB individuals, 117 CEU individuals, 115 YRI individuals). Briefly, the MDS analysis was performed with the filtered autosomal SNPs that were presented in both donors and Hapmap populations. The 20 samples were clustered with the Hapmap populations based on the MDS analysis, which is consistent with the reported ethnicity of these samples, including 16 Caucasian, 3 Hispanic, and 1 Asian (Supplementary Table 1, Supplementary Fig. 2a).

### Phasing with reference panel

The SNPs aligned between the 1000 genome phase 3 reference panel and the genomes of the 20 samples were extracted with shapeit2[57,58]. For each autosome, the overlapped SNPs of the sample genomes were phased with shapeit2 using the reference panel haplotypes with the same ethnicity as the sample group.

### snRNA-seq processing

The snRNA-seq raw data were processed with cell ranger. To remove the ambient RNA contamination, the read count matrix (gene x cell) was corrected with SoupX for each sample[59]. For each sample, to remove low quality cells, the corrected count matrix was filtered using the following parameters: min.cells = 5, nFeature_RNA ≥ 500, percent.mt ≤ 15 by Seurat[60]. To remove doublets, DoubletFinder was applied to each sample with doublet rate set at *the cell number*/1000 × 0.01[61]. After removing doublets for each sample, cell types were predicted using scPred based on the reference retinal cell atlas[28,62]. The expression of marker genes per cell type per sample were examined to confirm cell type assignment.

### snRNA-seq gene expression quantification

For each cell type, the average CPM of each gene across the cells from the same cell type of a sample was computed as the gene expression measurement per sample. For each cell type, the gene expression of all genes in the 20 samples were collected (gene x sample matrix) to perform quantile normalization. For each gene per cell type, the normalized gene expression levels were transformed using rank based inverse normal transformation[63]. For each cell type, only the genes with mean CPM (in the 20 samples) ≥5 were kept for downstream sc-eQTL analysis. To remove the effects of confounding variables (*e*.*g*., batch effect) from gene expression, the PEER factors were calculated from the transformed gene expression with the “PEER” R package[64,65].

### snATAC-seq processing

The snATAC-seq raw data were processed with cell ranger and then analyzed with ArchR[66]. The QC and filtering of low quality cells and doublets were performed with ArchR using the default setting (minTSS = 4 and minFrags = 1000, doublet filterRatio=1). The cell types of snATAC-seq were assigned by integrating the snRNA-seq data of the 20 samples using ArchR. For each sample, the snATAC-seq bam file per cell type per donor was generated according to the cell type label. For each cell type, the bam files from the same cell type of the 20 donors were merged to call snATAC peaks with macs3 in the default setting[67]. To reduce false positive peaks, only the peaks with mean FPKM ≥2 across samples per cell type were kept for each cell type. The filtered peaks from all cell types were combined to generate a set of standardized peak coordinates that can be compared among different cell types using the “Reduce” function in R. The peaks in the hg19 blacklist regions (wgEncodeHg19ConsensusSignalArtifactRegions) and chrY were filtered out. The standardized peak set was input into ArchR to generate peak to gene connection list, peak co-accessibility list, and the differential accessibility regions (DARs). The TFs were identified from the OCRs per cell type by chromVAR and correlating the TF expression with their motif enrichment across cell types (p.adj < 0.01, correlation coefficient > 0.5, and a maximum inter-cluster difference in deviation z-score > 75th percentile) with ArchR.

### sc-eQTL mapping

For each cell type, cis-eQTLs were mapped for the genes with mean CPM ≥5 using FastQTL[68]. Only the variants passing the following criteria were considered: 1) within +/-250 kb of gene TSS, 2) in OCRs of the given cell type, 3) with minor allele frequency (MAF) ≥0.1 across the 20 samples, and 4) minimum number of samples carrying the minor allele ≥ 4. Given the small sample size (N=20), three PEER factors and the first component of MDS analysis of the genotypes were used as covariates. The FastQTL were run in a nominal pass mode. To identify gene level significant sc-eQTLs, the p-value of each sc-eQTL per gene was corrected for multiple testing with Bonferroni method, based on the number of independent variants per gene estimated by eigenMT[69], for each cell type respectively.

### sc-ASE mapping

The snRNA-seq bam file per cell type per sample were generated according to the cell type label. To correct read mapping bias, the snRNA-seq bam file per cell type per sample were processed with WASP[70]. Duplicate reads were removed with UMI-tools[71]. For each sample, the reference-panel phased SNP VCF and corrected snRNA-seq bam files were used to generate haplotype count and genome-wide phased VCF with phASER[72]. The gene level haplotype counts for allelic expression were obtained using phASER Gene AE. For each cell type, the gene-level haplotypic counts per sample were combined to produce a haplotypic expression matrix (gene x sample) using phaser_expr_matrix.py of phASER-POP[73]. For each cell type, the effect sizes of all tested variant-gene pairs in the aforementioned sc-eQTL analysis were calculated using the aggregated haplotypic expression matrix and the genome-wide phased VCF with phaser_cis_var.py of phASER-POP. Only the variants with ≥4 heterozygotes are considered. For each cell type, genome-wide multiple testing correction was performed for each variant with Benjamini-Hochberg method. The variants with FDR <10% were identified as sc-ASEs.

### sc-ASCA mapping

For each sample, to correct read mapping bias, the snATAC-seq bam file per cell type per sample were processed with WASP[70]. Duplicate reads were removed with MarkDuplicates (Picard) from GATK[54]. The allelic count of SNPs was obtained using ASEReadCounter from GATK. For each SNP per cell type per sample with at least 10 reads from WGS and 10 reads from snATAC-seq are considered, and one-sided Fisher test was used to compare whether the allelic count ratio from snATAC-seq was significantly greater or less than the allelic count ratio of from WGS. For each cell type, the Fisher test P values of the same SNP in all heterozygous samples were combined to calculate a meta P value using the Stouffer’s method with the “metap” R package[74] (with the total read count in WGS-seq and snATAC-seq as the weight for each sample). For each SNP per cell type, only the meta P value in the effect direction with more significance was kept. For each cell type, the SNPs passing the follow criteria were considered: 1) with at least of three heterozygous samples and 2) considered in the aforementioned sc-eQTL analysis. To correct for genome-wide multiple testing, for each cell type, Benjamini-Hochberg correction was applied to meta P value of each SNPs to identify sc-ASCAs with FDR <10%.

### sc-caQTL mapping

For each cell type, the fragment count matrices (peak x sample) were generated based on the standardized peak coordinates in the given cell type and the snATAC-seq bam file (after WASP correction and removal of duplicate reads) per sample per cell type using featuerCounts[75]. For each cell type, the reference-panel phased SNPs were annotated with allelic read counts using RASQUAL tools[76]. To correct for library size and GC content bias in feature-level fragment counts per sample, the sample specific offset was computed using the rasqualCalculateSampleOffsets() function with the “GC-correction” option. The fragment count covariates were calculated with make_covariates() function of rasqual package (with variable number of covariates in different cell types) and were included into the model. For each cell type, sc-caQTL analysis was performed for the variants that were considered in sc-eQTL analysis. RASQUAL was run in two modes: 1) in the default setting and 2) with permuted sample labels using the “—random-permutation” option. To correct for multiple testing in feature level, the number of independent variants/tests per peak was determined with eigenMT[69].

Based on the number of independent tests, the true association P values and empirical permuted P values were corrected with Bonferroni method respectively. To correct for multiple testing genome-wide, the corrected true association P values were compared to the corrected empirical null distribution to determine the true P value threshold with FDR < 10%.

### LCRE identification

The gene-peak links were identified based on the correlation of gene expression and chromatin accessibility of snATAC-seq OCRs (-/+250kb) using the addPeak2GeneLinks() function in ArchR[66] with binarized peak read counts. The peak co-accessibility was estimated with the addCoAccessibility() function in ArchR with binarized peak read counts (for OCRs in -/+250kb). The snATAC-seq OCRs were annotated with ChIPseeker[77] and the OCRs within -/+1kb surrounding the promoter regions were defined as promoters. From the gene-peak links, we selected the OCRs that are not promoters as the CREs of the linked genes, while from the peak co-accessibility links, we selected the OCRs linked to promoters as the CREs of the target genes. The union set of gene-peak links and peak co-accessibility links were defined as the linked cis regulatory elements (LCRE) of the associated genes.

### Predicting the motif disrupting effects of SNPs

To determine if genetic variants within OCRs affect TF binding sites (TFBSs), we identified known TF motifs to the sequence surrounding genetic variants with motifBreakR[78], based on 2817 TF motifs (Hsapiens) from MotifBreakR database. The relative entropy of the motifs with reference allele and alternative allele was calculated, and only the TFBSs that were strongly affected (effect = “strong”) by SNPs were considered (with the parameters: filterp=TRUE, threshold= 1e-4, method=“ic”). We further required a TF with CPM ≥ 50 in the corresponding cell types to determine if its motif is perturbed by genetic variants.

### Identification of LD-independent sc-caQTL and LD-independent sc-eQTL

PLINK v1.90b5.2[55] (with the parameters: --clump-p1 0.05 --clump-p2 0.05 --clump-r2 0.50 --clump-kb 250) was used to clump sc-eQTLs per eGene per cell type and to clump sc-caQTLs per caPeak per cell type. The SNPs with the smallest p-value were assigned as the index SNPs. For multiple index SNPs with the same p-value, the SNP that is closest to gene TSS was assigned as the index sc-eQTL SNP, while the SNP that is closest to peak summit was assigned as the index sc-caQTL SNP.

### Identification of caQTLs associated with multiple genomic regions

For each common variant within snATAC-seq OCRs, we tested the association between the variant and the accessibility of snATAC-seq OCRs in -/+250kb surrounding the variant and took *p* < 0.005 as significant association. If the variant itself is a sc-caQTL of its residing OCR and also associated with other surrounding OCRs, we defined it as a sc-caQTL associated with multiple genomic regions and the residing OCR as the master caPeak while the other surrounding peaks as the dependent caPeaks. To avoid the confounding effect that two sc-caQTLs affecting two master caPeaks are in LD, the OCR that is a master caPeak and has its own resident caQTL that is in LD with the tested variant (*r*^2^ > 0.5) was filtered out.

### Cell type enrichment of GWAS loci

To determine the cell type enrichment of GWAS loci, we analyzed chromatin accessibility and gene expression derived from single cell multiomics data respectively. For chromatin accessibility, we partitioned the heritability of GWAS traits into the cell type OCRs and DARs through stratified LD score regression based on the summary statistics of GWAS traits with LDSC[43] (Supplementary Note). For gene expression, we assessed whether there is linear positive correlation between gene expression cell type specificity and gene-level genetic association with GWAS studies by MAGMA.Celltype[44–47] (Supplementary Note).

### Fine-mapping GWAS loci

We fine-mapped GWAS loci based on the summary statistics of GWAS studies[36,40,42,48]. For each GWAS study, the SNPs with *p* < 5 × 10^−8^ and present in 1000 genome (phase 3) European population were considered and were divided in the LD blocks identified by previous study[79]. The prior of each SNP was computed based on GWAS Z-score and the functional annotation of the SNP with “TORUS” package[80]. The annotation of a SNP was assigned to one of the categories: “4” if the SNP in the exonic/UTR regions, “3” if the SNP in the promoter region, “2” if the SNP in LCRE, “1” if the SNP in snATAC-seq OCR, “0” if the SNP not in snATAC-seq OCR. For each LD block, we calculated the PIP of each SNP and credible set of SNPs with the aforementioned prior weight generated by TORUS (i.e. functional PIP) and without the weighted prior (uniform PIP), respectively with “susieR” package[81]. Then we overlapped the fine-mapped variants with functional PIP > 0.1 with sc-eQTL, sc-caQTL and sc-ASCA.

## Declarations

### Ethics approval and consent to participate

All tissues were de-identified under HIPAA Privacy Rules. Institutional approval for the consent of patients for their tissue donation was obtained from the University of Utah and conformed to the tenets of the Declaration of Helsinki.

### Consent for publication

Not applicable.

### Data availability

The snRNA-seq and snATAC-seq data were deposited to latticeDB.

### Author contributions

RC, JW, XC, YL conceived the study and designed experiments. MD provide human samples. XC, YL, and LO performed the experiments. JW, QL and MW analyzed the data. JW, RC, XC and YL wrote the manuscript. All the authors edited and approved the manuscript.

## Acknowledgements

We acknowledge the computing cluster server in the Department of Molecular and Human Genetics and Human Genome Sequencing Center at Baylor College of Medicine for providing the computing resource.

## Additional Files

Supplementary Information: Supplementary Figures, Supplementary Table Titles and Supplementary Note.

Supplementary Table: Supplementary Tables 1-7

