## Supplementary Information for "Integrative single cell multiomics analysis of human retina indicates a role for hierarchical transcription factors collaboration in genetic effects on gene regulation"

### Supplementary Figures

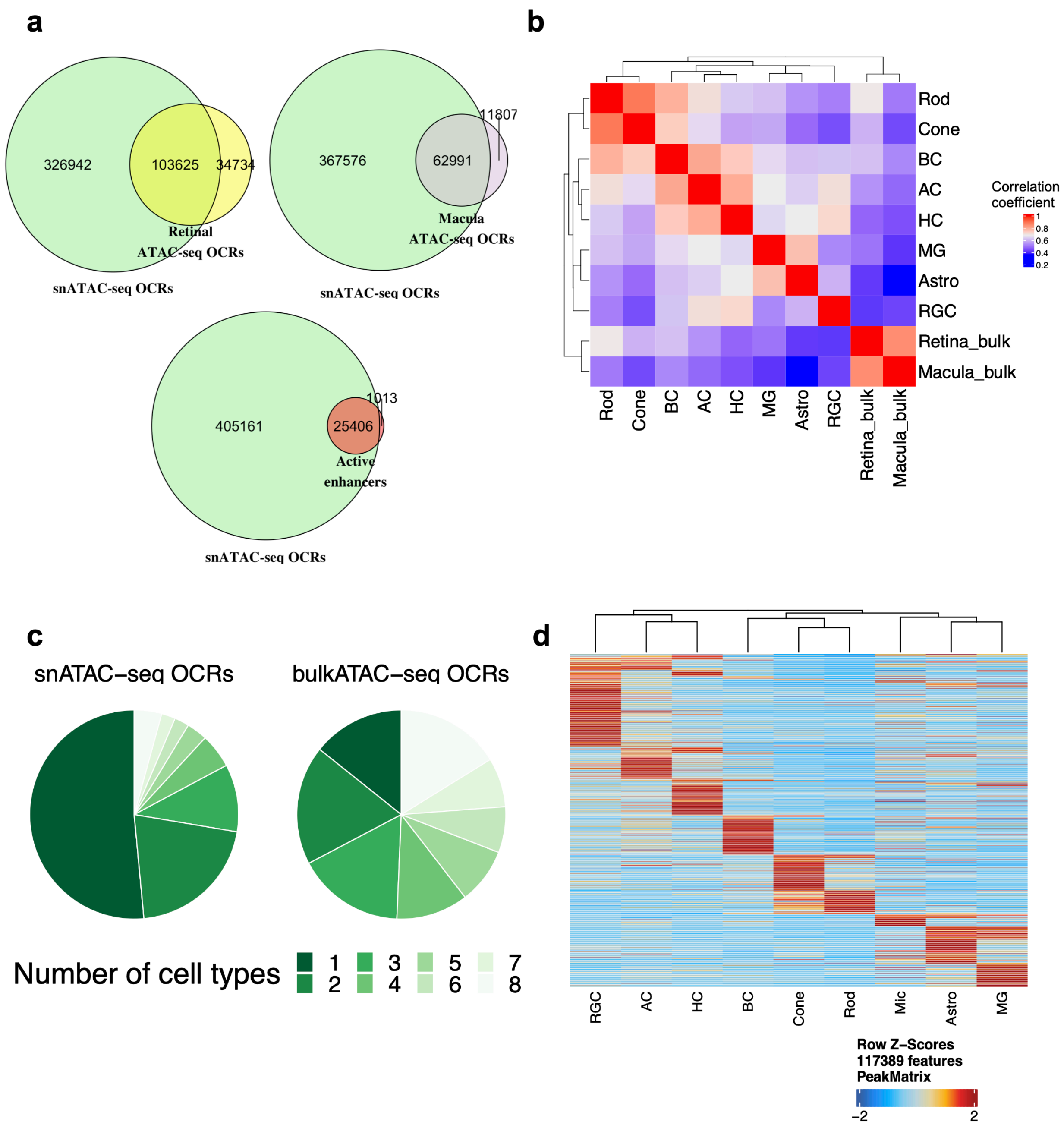


**Supplementary Fig. 1: The OCRs identified from snATAC-seq.**

**a,** The comparison of snATAC-seq OCRs with OCRs identified from bulk ATAC-seq and active enhancers identified from bulk ATAC-seq and ChIP-seq. **b,** The Pearson correlation of chromatin accessibility between retinal cell types and bulk retina and macula tissues. n=467806 OCRs. **c,** 51.5% of the OCRs from snATAC-seq are unique to one cell type while 14.3% of the OCRs from bulk ATAC-seq are unique to one cell type. **d,** Heatmap shows chromatin accessibility of differential accessible regions (DARs) in each retinal cell type. n = 117389 OCRs.

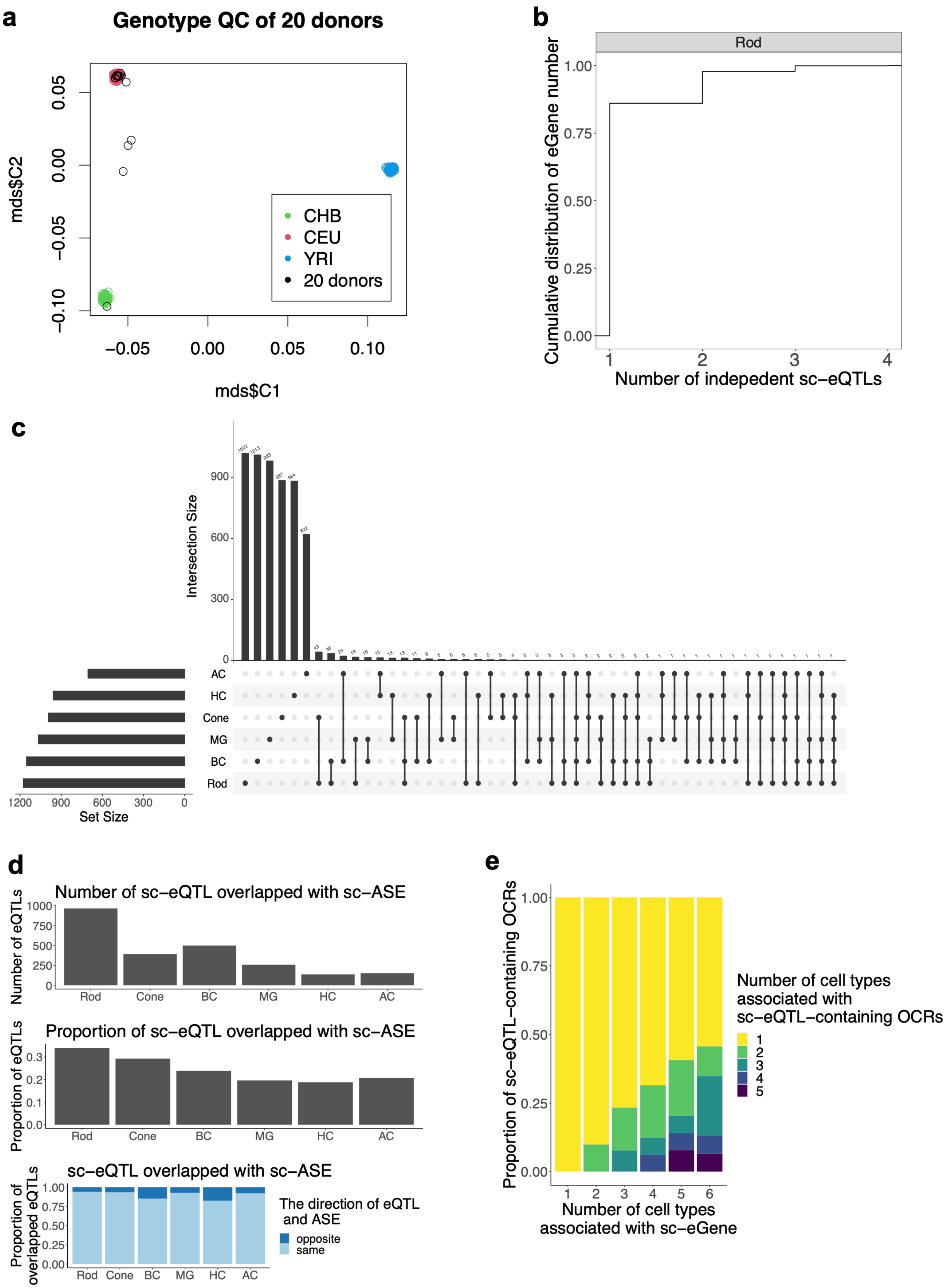
a
Genotype QC of 20 donors
mds$C2
mds$C1
0.05
0.00
−0.05
−0.10
−0.05
0.00
0.05
0.10
CHB
CEU
YRI
20 donors
b
Rod
Cumulative distribution of eGene number
1.00
0.75
0.50
0.25
0.00
1
2
3
4
Number of indepedent sc−eQTLs
c
Intersection Size
900
600
300
0
AC
HC
Cone
MG
BC
Rod
1200
900
600
300
0
Set Size
d
Number of sc−eQTL overlapped with sc−ASE
Number of eQTLs
1000
750
500
250
0
Rod
Cone
BC
MG
HC
AC
Proportion of sc−eQTL overlapped with sc−ASE
Proportion of eQTLs
0.3
0.2
0.1
0.0
sc−eQTL overlapped with sc−ASE
Proportion of overlapped eQTLs
1.00
0.75
0.50
0.25
0.00
The direction of eQTL and ASE
opposite
same
e
Proportion of sc−eQTL−containing OCRs
1.00
0.75
0.50
0.25
0.00
1
2
3
4
5
6
Number of cell types associated with sc−eGene
Number of cell types associated with sc−eQTL−containing OCRs
1
2
3
4
5

59

**Supplementary Fig. 2: Identification of sc-eQTL.**

**a,** MDS analysis of the genotype of the 20 donors and 316 individuals from the Hapmap populations. The X axis and Y axis indicate the first and second components from the MDS analysis. The green color indicates the Chinese Han population, the red color indicates the European population, the blue color indicates the African population. The 20 donor samples are colored in black. **b**, The majority of sc-eGenes have one independent sc-eQTL. The cumulative distribution of sc-eGene number shows the distribution of the number of independent sc-eQTLs per sc-eGene in Rod. n = 1011 sc-eQTLs. **c,** The number of sc-eQTLs across retinal cell types. **d,** The number and proportion of sc-eQTLs that are overlapped with sc-ASEs, based on the sc-eQTLs that were tested for sc-ASEs (top and middle). For the sc-eQTLs overlapped with sc-ASEs, the proportion of the variants with the same effect direction is colored in light blue, and the proportion with opposite effect direction is colored in dark blue (bottom). **e,** The OCRs with sc-eQTLs are different across cell types for the same sc-eGenes common in multiple cell types. n = 3447 OCRs for sc-eGenes associated with 1 cell type, n = 1368 for 2 cell types, n = 416 for 3 cell types, n = 162 for 4 cell types, n = 64 for 5 cell types, n = 46 for 6 cell types.

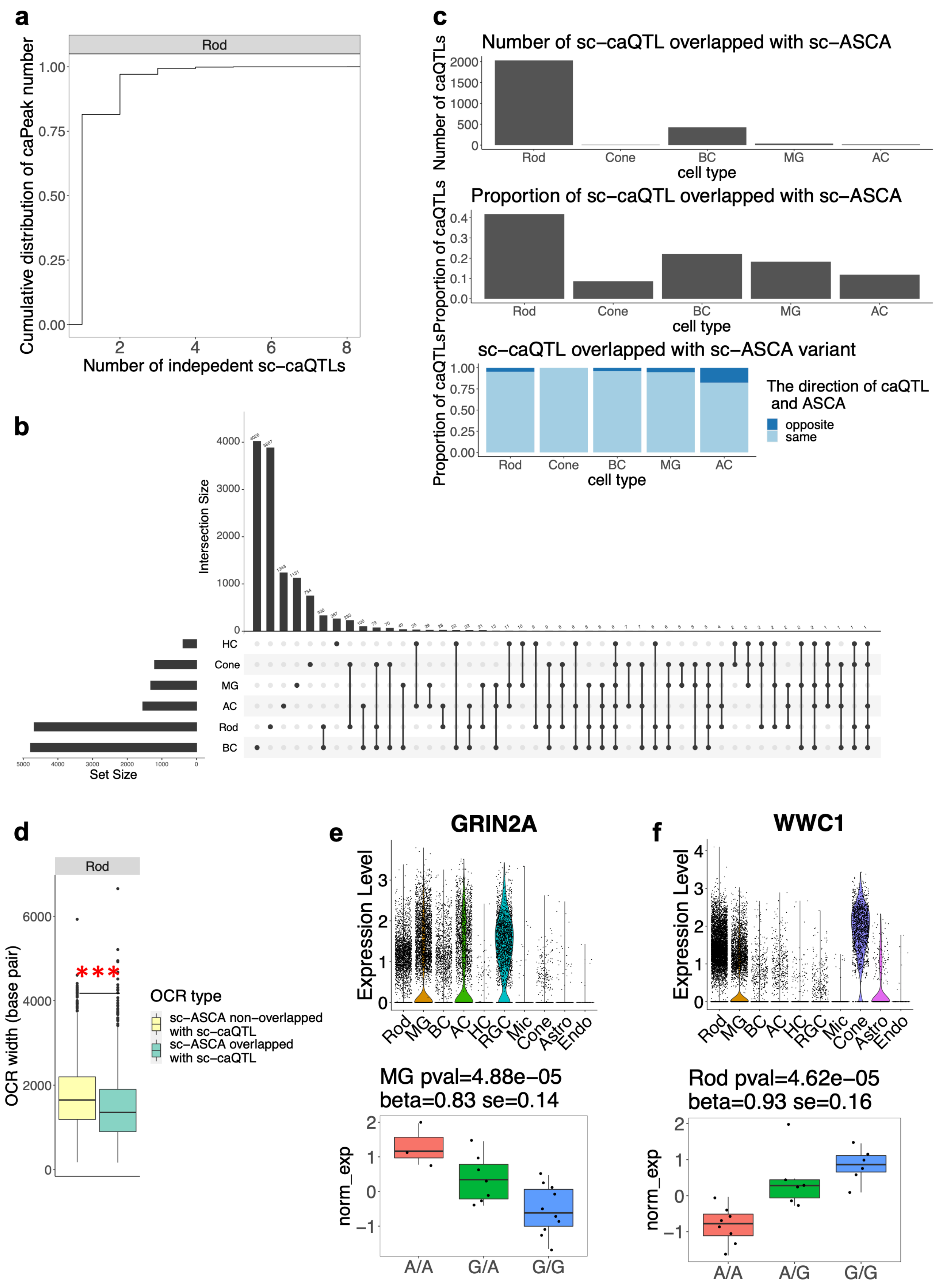

a
Rod
Cumulative distribution of caPeak number
1.00
0.75
0.50
0.25
0.00
2
4
6
8
Number of indepedent sc–caQTLs
c
Number of sc–caQTL overlapped with sc–ASCA
Number of caQTLs
2000
1500
1000
500
0
Rod
Cone
BC
MG
AC
cell type
Proportion of sc–caQTL overlapped with sc–ASCA
Proportion of caQTLs
0.4
0.3
0.2
0.1
0.0
Rod
Cone
BC
MG
AC
cell type
sc–caQTL overlapped with sc–ASCA variant
Proportion of caQTLs
1.00
0.75
0.50
0.25
0.00
Rod
Cone
BC
MG
AC
cell type
The direction of caQTL and ASCA
opposite
same
b
Intersection Size
4000
3000
2000
1000
0
HC
Cone
MG
AC
Rod
BC
5000
4000
3000
2000
1000
0
Set Size
d
Rod
OCR width (base pair)
6000
4000
2000
0
***
OCR type
sc–ASCA non–overlapped with sc–caQTL
sc–ASCA overlapped with sc–caQTL
e
GRIN2A
Expression Level
3
2
1
0
Rod
MG
BC
AC
HC
RGC
Mic
Cone
Astro
Endo
MG pval=4.88e−05
beta=0.83 se=0.14
norm_exp
2
1
0
−1
A/A
G/A
G/G
f
WWC1
Expression Level
4
3
2
1
0
Rod
MG
BC
AC
HC
RGC
Mic
Cone
Astro
Endo
Rod pval=4.62e−05
beta=0.93 se=0.16
norm_exp
2
1
0
−1
A/A
A/G
G/G

**Supplementary Fig. 3: Identification of sc-caQTL.**

**a,** The majority of sc-eGenes have one independent sc-caQTL. The cumulative distribution of OCR number shows the distribution of the number of independent sc-caQTLs per OCR in Rod. n = 3833 sc-caQTLs. **b,** The number of sc-caQTLs across cell types. **c,** The number and proportion of sc-caQTLs that are overlapped with sc-ASCAs, based on the sc-caQTLs that were tested for sc-ASCAs (top and middle). For the sc-caQTLs overlapped with sc-ASCAs, the proportion of the variants with the same effect direction is colored in light blue, and the proportion with opposite effect direction is colored in dark blue (bottom). **d,** The OCRs that contain sc-ASCA not overlapped with sc-caQTL are significantly wider than the OCRs that contain sc-ASCA overlapped with sc-caQTL in Rod. n = 1528 sc-ASCAs that are not overlapped with sc-caQTLs. n = 1393 sc-ASCAs that are overlapped with sc-caQTLs. One-sided Wilcoxon ranking sum test, $p < 1.47 \times 10^{-21}$. **e,** The sc-caQTL rs12447029 is also a sc-eQTL of GRIN2A in MG. **f**, The sc-caQTL rs6859300 is also a sc-eQTL of WWC1 in Rod.

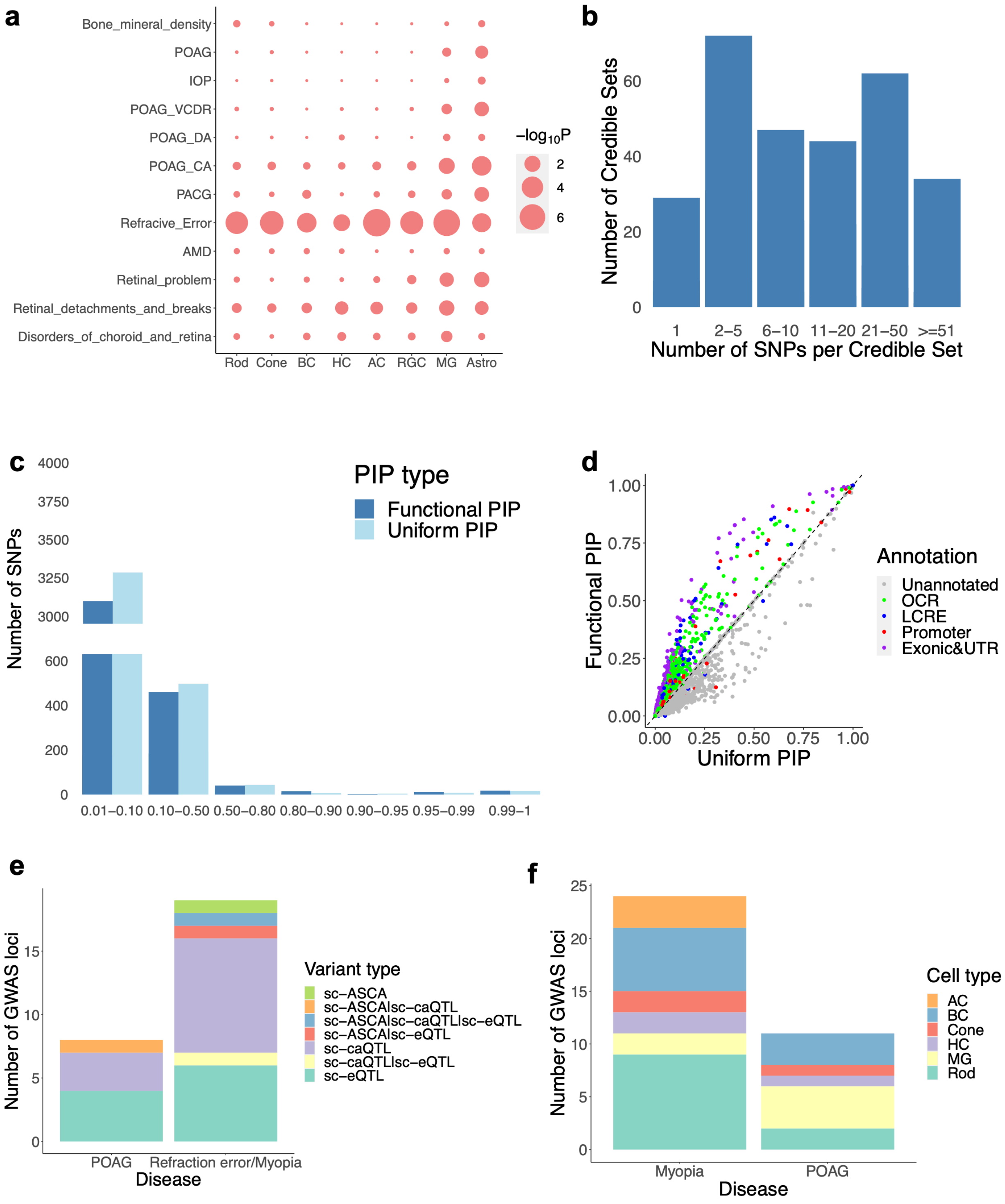


**Supplementary Fig. 4: Cell type enrichment and fine-mapping of GWAS loci.**

**a,** The cell type enrichment of 11 eye-related and one control GWAS loci by partitioning the heritability enrichment in cell type OCRs with LDSC. POAG: primary open-angle glaucoma. IOP:

intraocular pressure. VCDR: vertical cup-disc ratio of optic nerve. CA: cup area of optic nerve. DA: disc area of optic nerve. PCAG: primary angle closure glaucoma. AMD: age-related macular degeneration. **b,** The distribution of SNP numbers per credible set. **c,** The PIP value distribution of fine-mapped SNPs. Functional PIP: PIP computed with functional annotation informed prior, Uniform PIP: PIP computed without functional annotation informed prior. **d,** The variants in regulatory regions were prioritized with functional annotation informed prior (functional PIP), compared to the uniform PIP. n = 52087 SNPs. **e,** The sc-QTL and sc-ASCA distribution of the prioritized GWAS variants (PIP > 0.1). **f**, The cell type distribution of the prioritized GWAS variants overlapped with sc-QTL/ASCA.

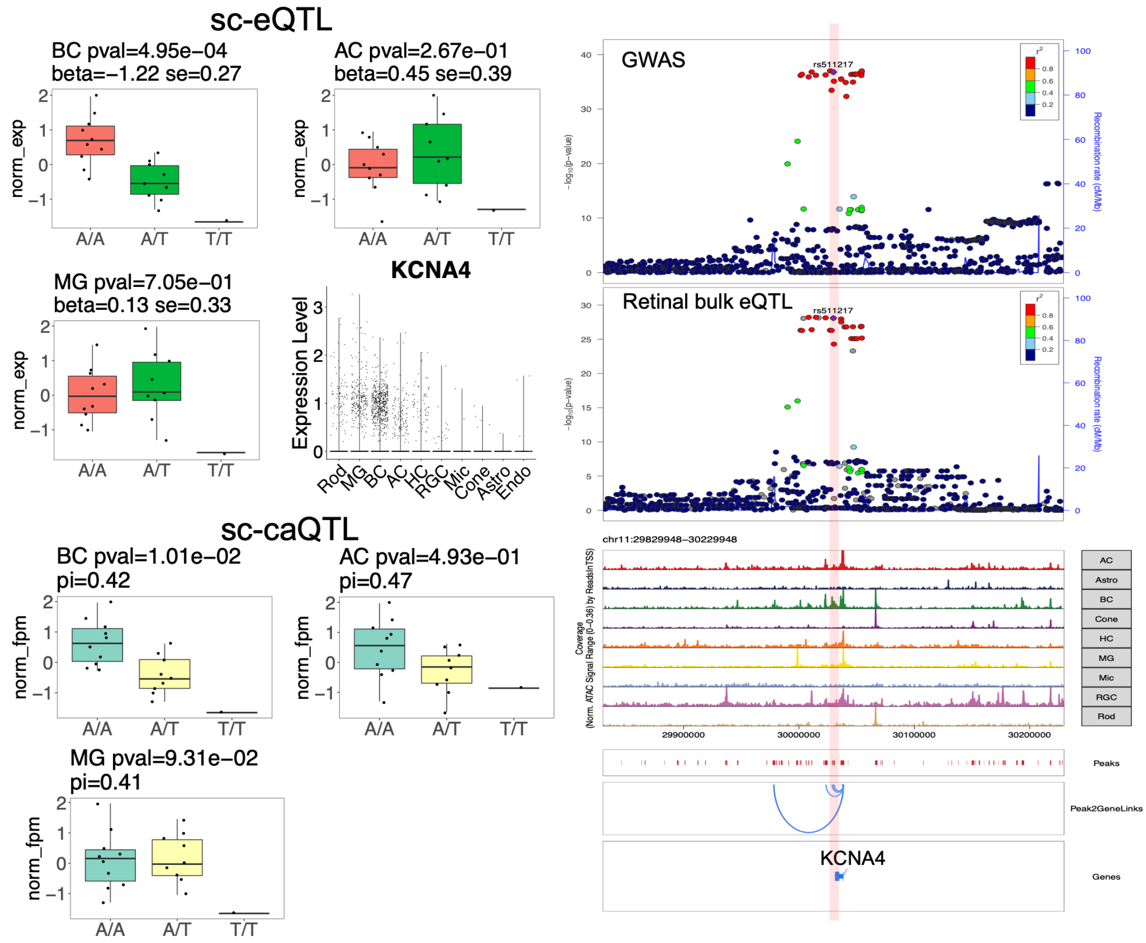


**Supplementary Fig. 5: An example of the prioritized GWAS candidate variants with the retinal bulk eQTL signal of its target gene colocalized with GWAS signal.**

The variant rs511217 associated with refraction error with PIP= 0.176 is a sc-eQTL of KCNA4 and a nominal significant sc-caQTL of its residing OCR in BC. This OCR is accessible in BC, MG and AC, and is a predicted LCRE of KCNA4. The retinal bulk eQTL signal of KCNA4 is also colocalized with the GWAS signal.

#### Supplementary Tables

**Supplementary Table 1.** The donor retina information.

**Supplementary Table 2.** The cell number per cell type per donor from snRNA-seq and snATAC-seq.

**Supplementary Table 3.** snATAC-seq OCR list.

**Supplementary Table 4.** The summary statistics of sc-eQTL with gene level FDR < 0.1 per cell type.

**Supplementary Table 5.** The summary statistics of sc-caQTL with genome level FDR < 0.1 per cell type.

**Supplementary Table 6.** The fine-mapped GWAS variants with PIP > 0.1 and overlapped with sc-QTL and/or sc-ASCA.

**Supplementary Table 7.** The target genes of the prioritized GWAS variants with PIP > 0.1 and overlapped with sc-QTL and/or sc-ASCA.

### Supplementary Note

#### The QC and filtering of the WGS variants

QC and filtering of the WGS variants were performed with the following criteria: 1) Indicated as "PASS" by GATK based on the filtering criteria: For SNP: QD < 2.0, QUAL < 30.0, SQR > 3.0, FS > 60, MQ < 40, MQRankSum < -12.5, ReadPosRankSum < -8.0. For INDEL: QD < 2.0, QUAL < 30.0, FS > 200, ReadPosRankSum < -20.0. 2) Remained dimorphic after assigning the variants as missing with the following criteria: a) the variants not called; b) variant calls with allelic imbalance (AB>0.9 or AB<0.1); c) variant GQ per sample (from GATK) < 20; d) heterozygous genotype calls on chrX nonPAR of the male donors; e) heterozygous genotype calls on chrY nonPAR of the male donors or, heterozygous/homozygous calls on chrY nonPAR of female donors. 3) A missingness rate ≤ 20%. 4) Genomic sites not involving MNP. 5) Not in low complex regions. 6) Passing Hardy-Weinberg Equilibrium testing. The QC and filtering were performed using GATK, plink, bcftools and custom scripts in Perl[1,2].

#### Processing of the bulk ATAC-seq and ChIP-seq data

The human retina and macula bulk ATAC-seq and bulk ChIP-seq of H3K27ac and H3K4me2 data were downloaded from https://www.ncbi.nlm.nih.gov/geo/query/acc.cgi?acc=GSE13731129[3]. The sequencing reads were trimmed with Trimmomatic-0.39 and mapped to the hg19 genome assembly with bowtie2-2.3.5.1 (The alignment bam files were sorted by samtools-1.2)[4–6]. The narrowPeaks were called from the bam files with Genrich (https://github.com/jsh58/Genrich). The narrowPeaks from the bulk ATAC-seq and ChIP-seq were then intersected with snATAC-peaks with bedtools[7].

#### The colocalization of GWAS loci with retina bulk eQTL

We identified SNPs with $p < 10^{-6}$, focused on -/+500kb surrounding the SNP, and considered the genes whose TSS in the region. Then we extracted the GWAS variants and retinal bulk eQTL variants[8] of the genes in the region and performed colocalization using the "coloc" R package (v5.1.0) with default parameter setting based on the summary statistics of the GWAS study and retinal bulk eQTLs for each of the corresponding genes respectively[9,10]. We identified the genes with the posterior probabilities of colocalization (PP4) > 0.5 as the genes colocalized with GWAS signals[11].

**Identification of cell type enrichment underlying GWAS traits**

For chromatin accessibility, we partitioned the heritability of GWAS traits into the cell type OCRs and DARs through stratified LD score regression based on the summary statistics of GWAS traits with LDSC[12]. First, for each GWAS study, the SNPs that are overlapped with HapMap3 SNPs were annotated based on whether they are in the OCRs or DARs in each cell type. Then, LD-scores of these SNPs were calculated within 1 cM windows based on the 1000 Genome data. For each GWAS study, the LD-scores of these SNPs were integrated with the ones from baseline model, which include non-cell type specific annotation, downloaded from https://alkesgroup.broadinstitute.org/LDSCORE/. Finally, for each GWAS study, the heritability in the annotated genomic regions were estimated and compared with the baseline model to determine if regions in each cell type were enriched with the heritability of the corresponding GWAS trait.

For gene expression, we assessed whether there is linear positive correlation between gene expression cell type specificity and gene-level genetic association with GWAS studies by MAGMA.Celltype[13–16]. We formatted GWAS summary statistics with the "MungeSumstats" R package, included SNPs -35kb/+10kb of each gene based on 1000 genome "eur" population, formatted snRNA-seq expression data with the "EWCE" R packages, and detected the linear enrichment with MAGMA.Celltype[13–16]. The enrichment p-value was further corrected with Benjamini-Hochberg method given the number of cell types and GWAS studies considered.
